# Separating Functions of the Phage-Encoded Quorum-Sensing-Activated Antirepressor Qtip

**DOI:** 10.1101/2019.12.16.877910

**Authors:** Justin E. Silpe, Andrew A. Bridges, Xiuliang Huang, Daniela R. Coronado, Olivia P. Duddy, Bonnie L. Bassler

## Abstract

Quorum sensing is a process of chemical communication that bacteria use to track cell density and coordinate gene expression across a population. Bacteria-infecting viruses, called phages, can encode quorum-sensing components that enable them to integrate host cell density information into the lysis-lysogeny decision. Vibriophage VP882 is one such phage, and activation of its quorum-sensing pathway leads to the production of an antirepressor called Qtip. Qtip interferes with the prophage repressor (cI_VP882_), leading to host-cell lysis. Here, we show that Qtip interacts with the N-terminus of cI_VP882_, inhibiting both cI_VP882_ DNA-binding and cI_VP882_ autoproteolysis. Qtip also sequesters cI_VP882_, localizing it to the poles. Qtip can localize to the poles independently of cI_VP882_. Alanine-scanning mutagenesis of Qtip shows that its localization and interference with cI_VP882_ activities are separable. Comparison of Qtip to a canonical phage antirepressor reveals that, despite both proteins interacting with their partner repressors, only Qtip drives polar localization.

## Introduction

Quorum sensing (QS) is a bacterial cell-cell communication process that controls collective behaviors. QS bacteria produce, release, and detect accumulated signaling molecules called autoinducers (AIs) [reviewed in (Papenfort and Bassler, 2016)]. QS-controlled behaviors include biofilm formation (Hammer and Bassler, 2003), virulence factor production (Miller et al., 2002; Zhu et al., 2002), competence (Meibom, 2005), and deployment of CRISPR-Cas defense systems that eliminate incoming foreign DNA from plasmids and bacteria-specific viruses called phages (Høyland-Kroghsbo et al., 2017). Phages also engage in chemical dialogs. For example, some *Bacillus* phages encode a phage-phage communication module termed the arbitrium system (Erez et al., 2017; Stokar-Avihail et al., 2019). In this case, phage-encoded proteins drive the production and release of signaling peptides, that, when detected by other prophages in the bacterial community, launch a lysogeny-activating program across the infected bacterial population. Phage-encoded QS components resembling host bacterial QS components have been identified using bioinformatics (Hargreaves et al., 2014; Silpe and Bassler, 2019a, 2019b), and in systems that have been tested, the phage-encoded QS receptors bind and respond to the AI molecule produced by the host bacterium. In one such instance, vibriophage VP882 encodes a QS receptor, called VqmA_Phage_, which is homologous to the bacterial QS VqmA receptor in *Vibrio cholerae* (Silpe and Bassler, 2019a). Both the host- and phage-encoded proteins bind to the host-produced AI, 3,5-dimethyl-pyrazin-2-ol (DPO), however, the AI bound VqmA proteins control different pathways. Host VqmA binding to DPO regulates *V. cholerae* QS behaviors (Herzog et al., 2019; Papenfort et al., 2017), while binding of DPO by the phage-encoded VqmA activates lytic development, resulting in killing of the *V. cholerae* host (Silpe and Bassler, 2019a). In a similar vein, recently, the QS AI called 4,5-dihydroxy-2,3-pentanedione (DPD), that is produced by many species of bacteria (Chen et al., 2002; Schauder et al., 2001), has been shown to promote lytic behavior in phage T1 of *Escherichia coli* (Laganenka et al., 2019). How phage T1 detects the DPD AI is not yet understood.

Phage lysis-lysogeny decisions are often controlled by phage proteins that repress expression of genes encoding activators of phage lytic genes. Relief of repression launches the phage lytic programs. Thus, the transition from lysogeny to lysis requires a mechanism to inactivate the phage repressor [reviewed in (Ptashne, 2004)]. Commonly, phage repressor proteins are subject to post-translational regulation. In the case of phage lambda, the cI repressor protein (cI_Lambda_) is auto-cleaved in an SOS-activated, RecA-dependent manner (Roberts and Roberts, 1975; Sauer et al., 1982). Cleavage abolishes cI_Lambda_ repressor activity. Repressors of other phages, for example, that of coliphage 186, are, by contrast, inactivated via protein-protein interactions with small antirepressor proteins (Shearwin et al., 1998). In these cases, production of the antirepressor is frequently controlled by a LexA-regulated promoter, and thus, despite different mechanisms, as in phage lambda, repressor inactivation is induced by the host SOS response (Heinzel et al., 1992; Kim et al., 2016; Lemire et al., 2011; Mardanov and Ravin, 2007; Quinones et al., 2005; Shearwin et al., 1998). Unlike autoproteolytic repressors, however, the mechanisms underlying antirepressor-mediated inactivation of phage repressors have seldom been fully characterized and it is possible that a given antirepressor can function by more than one mechanism. Known mechanisms include functioning as a competitive inhibitor of repressor DNA binding [reviewed in (Wang et al., 2014)], binding an allosteric site on a partner repressor protein and forcing it to adopt a DNA-incompetent confirmation (Kim et al., 2016), and sequestering the repressor rendering it insoluble (Davis et al., 2002).

The repressor encoded by vibriophage VP882, called cI_VP882_, can be inactivated by two mechanisms: RecA-dependent cleavage of the cI_VP882_ repressor occurs, similarly to lambda, following activation of the SOS response (Silpe and Bassler, 2019a). Additionally, antirepressor-mediated inactivation of the cI_VP882_ repressor is triggered by the phage-encoded QS pathway. Specifically, when VqmA_Phage_ binds the host-produced DPO AI, the complex activates transcription of a phage gene encoding an antirepressor, Qtip, that inactivates cI_VP882_ ((Silpe and Bassler, 2019a) and Figure 1A). By tuning into the SOS-response, VP882 prophages can connect their lysis-lysogeny decisions to the viability of the host and, by monitoring local bacterial population density, VP882 prophages can gauge the prospect of encountering future hosts.

**Figure 1:**
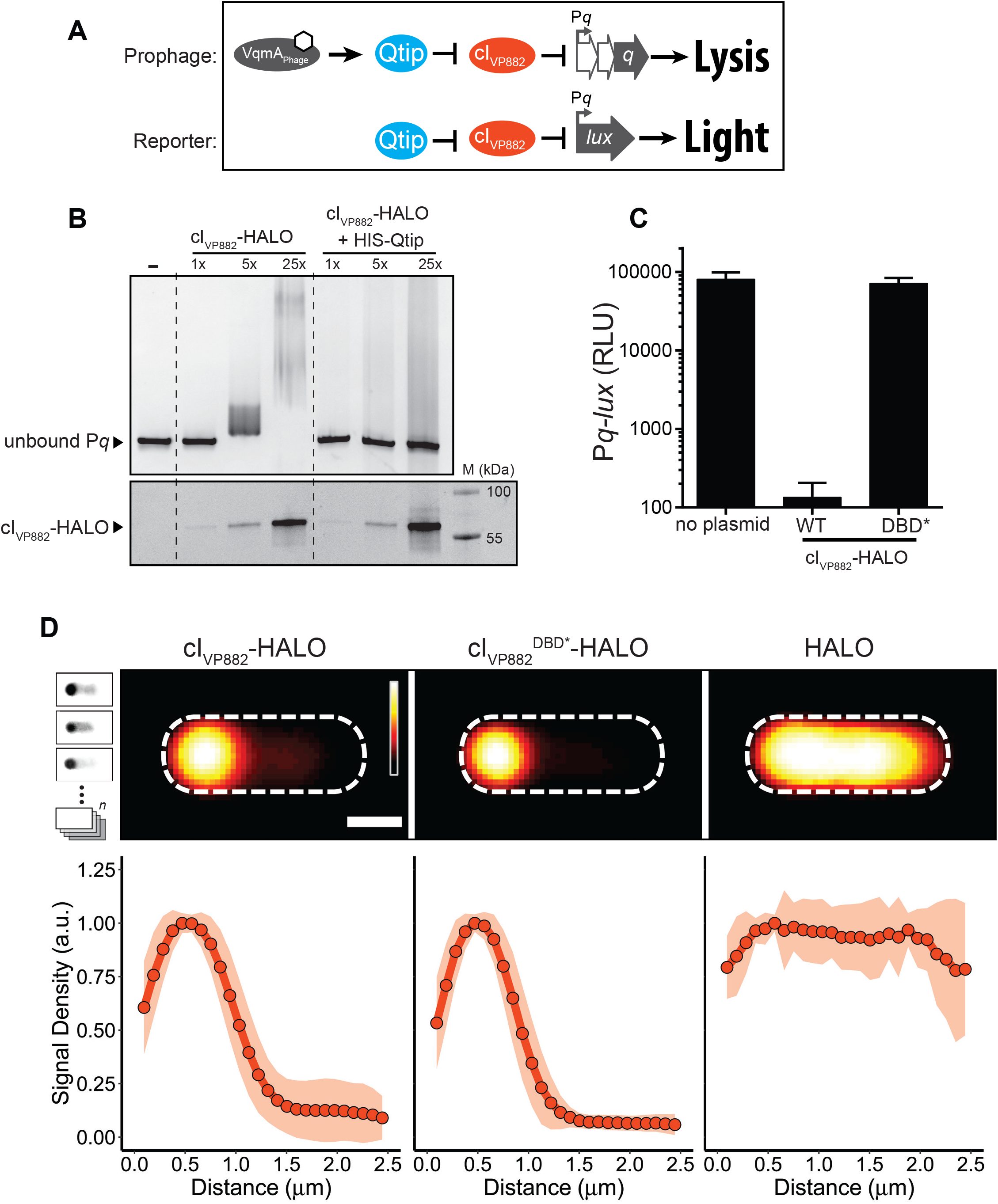
cI_VP882_ DNA-binding is blocked by Qtip but a cI_VP882_ mutant lacking DNA-binding capability does not prevent Qtip recognition. (A) Schematic depicting the phage VP882-encoded QS pathway and the reporter system used in this work. Top: VqmA_Phage_, when bound to DPO (white hexagon), activates expression of the *qtip* gene encoding the Qtip antirepressor. Qtip inactivates the cI_VP882_ repressor, enabling expression of P*q* and subsequent Q-mediated host cell lysis. Bottom: The reporter system used in this work to monitor Qtip and cI_VP882_ activity. P*q* is fused to the luciferase operon (*lux*). Light production is low when the cI_VP882_ repressor is functional and light production is high when Qtip is active and/or the cI_VP882_ repressor is non-functional. (B) Upper panel: EMSA analysis of P*q* DNA retarded by cI_VP882_-HALO protein purified alone or in complex with HIS-Qtip. The relative amount of cI_VP882_-HALO in each lane is indicated (1x = ~12 nM). Lower panel: The reactions from the upper panel were subjected to SDS-PAGE analysis and imaged using a Cy5 filter set to visualize HALO-Alexa_660_, to which the cI_VP882_-HALO protein had been conjugated. The molecular weight marker is designated M. (C) Light production from *E. coli* harboring the P*q-lux* reporter (no plasmid) or the P*q-lux* reporter and a plasmid encoding WT cI_VP882_ or the cI_VP882_^DBD*^-HALO allele. Relative light units (RLU) were calculated by dividing bioluminescence by OD_600_. Data represented as mean ± SD with n = 3 biological replicates. (D) Upper panel: Average individual cell images of recombinant *E. coli* producing Qtip and either, cI_VP882-_HALO, cI_VP882_^DBD*^-HALO, or the HALO tag. HALO-TMR fluorescence intensity is displayed as a red heat map (black and white reflect the lowest and highest intensity, respectively). Dashed lines denote the average cell outlines. Scale bar = 1 μm. Upper left inset: Schematic showing aligned individual cells averaged to produce composite images in the upper panel (n = 20-25 cells per condition, see Methods). In the schematic, the HALO-TMR fluorescence intensity from three representative individual cells harboring Qtip and cI_VP882_^DBD*^-HALO is shown in inverted greyscale. Lower panel: Line plots of HALO-TMR fluorescence intensity extracted from individual cell images used to generate the composite images displayed in the upper panel. The distance along the x-axis is relative to the left most edge of each cell. Shaded regions represent ± 1 SD from the mean.

Qtip sequesters cI_VP882_, and Qtip-cI_VP882_ aggregates are localized at the cell poles (Silpe and Bassler, 2019a). However, the mechanism by which Qtip inactivates the cI_VP882_ repressor remains unknown. Qtip is a small protein (~8 kDa) with no predicted domains and no close homologs, preventing structure-function and homology-based predictions. To probe the Qtip antirepressor mechanism, here, we developed *in vitro* assays to show that Qtip inhibits cI_VP882_ DNA binding and autoproteolysis. We demonstrate that, while the C-terminal domain of cI_VP882_ possesses the autocleavage activity, Qtip recognizes the N-terminal domain of cI_VP882_, which harbors the cI_VP882_ DNA binding activity. Mutation of some of the amino acid residues on cI_VP882_ that eliminate DNA binding do not interfere with Qtip binding to cI_VP882_, while other mutations eliminate both activities. Qtip inactivates repressors similar to cI_VP882_ but not those of lambda or phage P22. Likewise, Ant, the antirepressor from phage P22, inactivates repressors from lambda and P22 (Susskind and Botstein, 1975) but not from phage VP882. We show that, unlike Qtip, Ant does not alter the localization of its partner repressors. We mutagenized Qtip to identify residues essential for disabling cI_VP882_ DNA binding and residues required for polar localization. We show that these two Qtip properties are separable.

## Results

### Qtip Prevents cI_VP882_ from Binding to DNA

Previous work on phage VP882 showed that production of Qtip counteracts repression by cI_VP882_ at the target *q* promoter, leading to production of the Q antiterminator, launch of the lytic program, and host-cell lysis (Figure 1A and (Silpe and Bassler, 2019a)). Moreover, microscopy demonstrated that Qtip alters the localization of cI_VP882_. In the absence of Qtip, cI_VP882_ is diffuse in the cytoplasm, and when Qtip is present, cI_VP882_ forms foci colocalized with Qtip at the cell poles (Silpe and Bassler, 2019a).

We sought to determine how Qtip binding alters the ability of cI_VP882_ to function as a repressor. We first examined whether Qtip interferes with cI_VP882_ binding to DNA. To do this, we co-expressed N-terminal-hexahistidine-tagged Qtip (HIS-Qtip) with C-terminal-HALO-tagged cI_VP882_ (cI_VP882_-HALO) and using Ni-NTA resin, purified HIS-Qtip in complex with cI_VP882_-HALO. In parallel, we purified the cI_VP882_-HALO protein using heparin-affinity chromatography. We measured cI_VP882_-HALO binding and binding of the cI_VP882_-HALO-His-Qtip complex to the cI_VP882_ target promoter, called P*q*. Electromobility shift assays (EMSA) showed that the cI_VP882_-HALO protein shifted the P*q* DNA, however, the cI_VP882_-HALO protein purified in complex with HIS-Qtip failed to do so (Figure 1B). These results indicate that Qtip binding to cI_VP882_ prevents cI_VP882_ from binding target DNA, possibly providing the mechanism by which, in phage VP882 lysogens, Qtip initiates the lytic program.

We wondered whether the amino acid residues required for cI_VP882_ to bind DNA are also required for cI_VP882_ to interact with Qtip. We used domain analysis to identify 6 residues in the cI_VP882_ N-terminus that are predicted to participate in sequence-specific DNA-binding (InterProScan): L18, Q19, G32, S35, R39, and G40. Pairing these sites with our knowledge of cI homologs that Qtip does and does not sequester (Silpe and Bassler, 2019a), we aligned the Qtip-interacting cI proteins and found that 2 of these 6 sites (Q19 and S35) are shared between all of the susceptible proteins and are absent in the analogous positions in cI_Lambda_ (Figure S1A), which is not sequestered by Qtip (Silpe and Bassler, 2019a). We mutated cI_VP882_ to make cI_VP882_^Q19A/S35A^ and fused the double mutant protein to HALO. We call this construct cI_VP882_^DBD*^-HALO. We tested the ability of cI_VP882_^DBD*^-HALO to repress its target P*q* promoter using recombinant *E. coli* harboring a plasmid carrying the *q* promoter fused to the luciferase operon (P*q*-*lux*). WT cI_VP882_-HALO repressed P*q-lux* expression while cI_VP882_^DBD*^-HALO did not (Figure 1C), indicating that the Q19A/S35A double mutation leads to a DNA binding defective cI_VP882_. We also assessed whether Qtip could sequester cI_VP882_^DBD*^-HALO by performing confocal microscopy on recombinant *E. coli* carrying Qtip and either WT cI_VP882_-HALO, cI_VP882_^DBD*^-HALO, or only HALO (no cI_VP882_), each labeled with HALO-TMR. The upper panel in Figure 1D shows composite images of the average HALO-TMR fluorescence signals from aligned individual cells expressing the specified constructs. We also measured the fluorescence intensity across the long axis of the aligned individual cells, displayed as an average line profile of fluorescence intensity as a function of the distance from the left cell pole as depicted in the figure (lower panel). We found that Qtip sequestration of cI_VP882_^DBD*^-HALO was indistinguishable from Qtip sequestration of WT cI_VP882_^DBD*^-HALO (Figure 1D), indicating that the residues required for the cI_VP882_ repressor to bind DNA are not required for Qtip to bind to the cI_VP882_ repressor.

### RecA* ctivates and Qtip Inhibits cI_VP882_ Autoproteolysis

In addition to binding DNA and Qtip, the cI_VP882_ repressor is subject to RecA-dependent autocleavage (Silpe and Bassler, 2019a). Based on domain prediction and homology modeling using the cI_Lambda_ repressor and other LexA-like repressor proteins as references, we predicted that the cI_VP882_ N-terminal domain is responsible for DNA binding while cleavage is enabled by a catalytic domain (S24 peptidase-like) residing in the C-terminus. To assess autoproteolysis, we established *in vitro* conditions under which cI_VP882_ cleavage could be monitored. We incubated purified cI_VP882_-HALO protein with single stranded DNA (ssDNA), RecA, and ATP-γ-S. ssDNA and ATP-γ-S are known to stimulate RecA to adopt an activated state (termed RecA*), which, in the presence of other repressor proteins (e.g., LexA and cI_Lambda_) promotes their autoproteolytic activity (Giese et al., 2008; Ndjonka and Bell, 2006). Figure 2A shows that under these conditions, cI_VP882_-HALO cleavage occurred within 10 min at 37°C, while in reactions lacking ATP-γ-S or RecA, cI_VP882_-HALO did not undergo cleavage. The data in Figure 1B show that Qtip inhibits cI_VP882_ DNA binding. We examined whether Qtip could also affect cI_VP882_ autoproteolysis. Figure 2B shows that, in contrast to when cI_VP882_-HALO was purified alone, cI_VP882_-HALO purified in complex with Qtip did not undergo cleavage, suggesting that Qtip binding inhibits cI_VP882_ autoproteolytic activity.

**Figure 2:**
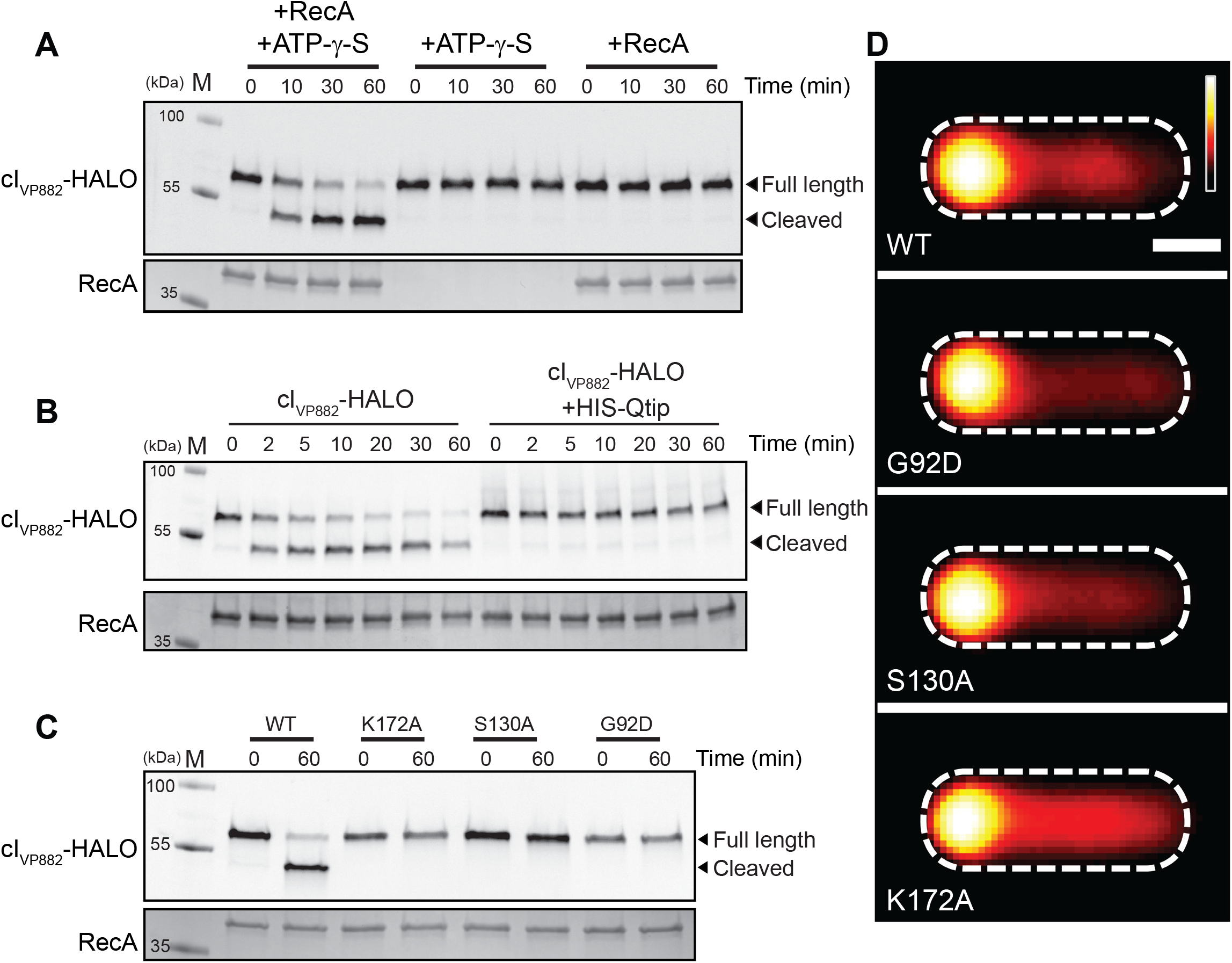
Qtip prevents *in vitro* cleavage of cI_VP882_ but cI_VP882_ mutants lacking cleavage or catalytic sites do not prevent Qitp recognition. (A) *In vitro* cleavage of cI_VP882_-HALO monitored by SDS-PAGE analysis. The presence of RecA and/or ATP-γ-S in the reactions is indicated above each lane. (B) *In vitro* cleavage of cI_VP882_-HALO purified alone or in complex with HIS-Qtip. Samples were concentration matched according to the cI_VP882_-HALO Alexa_660_ signal. (C) *In vitro* cleavage of WT cI_VP882_-HALO and the indicated catalytic-site (K172A and S130A) and cleavage-site (G92D) variants. For (A-C): all cI_VP882_-HALO proteins were labeled with HALO-Alexa_660_. Upper panels, gels imaged using a Cy5 filter set for HALO-Alexa_660_ detection; lower panels, the same gels stained for total protein. Incubation times noted above each lane. Marker, M. In (B) and (C), all samples contained ATP-γ-S and RecA. (D) Composite images from individual cell analyses of *E. coli* producing Qtip and either WT cI_VP882_-HALO or the cleavage-site (G92D) or catalytic-site (S130A and K172A) cI_VP882_-HALO variant. HALO-TMR fluorescence intensity and scale bar represented as in Figure 1D.

We investigated the requirements for Qtip inhibition of cI_VP882_ autoproteolytic activity. Based on homology to cI_Lambda_, we predicted that cleavage of cI_VP882_ occurs between residues A91 and G92 (Figure S1A), analogous to the cleavage site between A111 and G112 in cI_Lambda_ (Sauer et al., 1982), and that the two active site residues are S130 and K172 (Figure S1A), which match the known catalytic residues S149 and K192 in cI_Lambda_ (Slilaty and Little, 1987). Consistent with these predictions, Figure 2C shows that the cI_VP882_^S130A^-HALO and cI_VP882_^K172A^-HALO catalytic site mutants and the cI_VP882_^G92D^-HALO cleavage site mutant did not undergo autoproteolysis. Importantly, however, Figure 2D shows that, like WT cI_VP882_-HALO, the cI_VP882_-HALO cleavage site and catalytic site mutants could all still be sequestered by Qtip. Thus, Qtip binding prevents cI_VP882_ autoproteolysis, but the residues involved in cI_VP882_ cleavage and catalysis are not required for Qtip to do so.

### cI_VP882_ Autoproteolysis Occurs by an Intra-Molecular Mechanism

To probe the mechanism of cI_VP882_ autoproteolysis, we relied on established mechanisms underlying self-cleavage of related repressor proteins. Specifically, cleavage can occur via an intra-molecular mechanism, as in the case of LexA and cI_Lambda_ (Slilaty et al., 1986), or an inter-molecular mechanism, as in the case of UmuD of *E. coli* (McDonald et al., 1998). To classify the catalytic behavior of cI_VP882_, we combined WT cI_VP882_-HALO with each of the catalytic-site variants, cI_VP882_^S130A^-HALO or cI_VP882_^K172A^-HALO. Our rationale was that if cI_VP882_ is cleaved by an inter-molecular mechanism, then WT cI_VP882_-HALO, in addition to cleaving other WT cI_VP882_-HALO proteins, would cleave the catalytically dead cI_VP882_^S130A^-HALO and cI_VP882_^K172A^-HALO mutants *in trans*. By contrast, if intra-molecular cleavage is the mechanism, only the WT cI_VP882_-HALO protein would be cleaved.

To distinguish WT cI_VP882_-HALO from the cI_VP882_-HALO catalytic site mutants in our mixed reactions, we took advantage of the fact that the HALO tag can be labeled with different fluorophores and the fluorophores remain covalently bound and continue to fluoresce when excited under denaturing SDS-PAGE conditions (Figure S1B). Thus, we employed a red-fluorescent ligand, HALO-TMR, to label each of the catalytic site mutants (cI_VP882_^S130A^-HALO and cI_VP882_^K172A^-HALO) and a far-red/IR ligand, HALO-Alexa_660_, to label WT cI_VP882_-HALO. In our mixed reactions, the differentially tagged proteins could be resolved using filter sets that are specific to the excitation-emission spectrum of each fluorophore. The overlaid emission output of cI_VP882_-HALO conjugated to different fluorophores, prior to-and after mixing, is shown in Figure S1B. Mixing of the far-red labeled WT cI_VP882_-HALO with either of the red-fluorescent-labeled catalytic site mutants shows that only the WT (Alexa_660_-labeled) protein was cleaved (Figure 3A), indicating that the cI_VP882_ cleavage mechanism is intra-molecular. To eliminate the possibility that the red-fluorescent HALO ligand prevents cleavage of cI_VP882_-HALO, we exchanged the fluorophores used to label the different proteins. Again, consistent with an intra-molecular cleavage mechanism, only the WT (TMR-labeled) cI_VP882_-HALO protein was cleaved (Figure 3A). We conclude that cI_VP882_ is cleaved likely by the same intra-molecular mechanism as cI_Lambda_, and cleavage relies on conserved catalytic residues and the identical alanyl-glycyl cleavage sequence.

**Figure 3:**
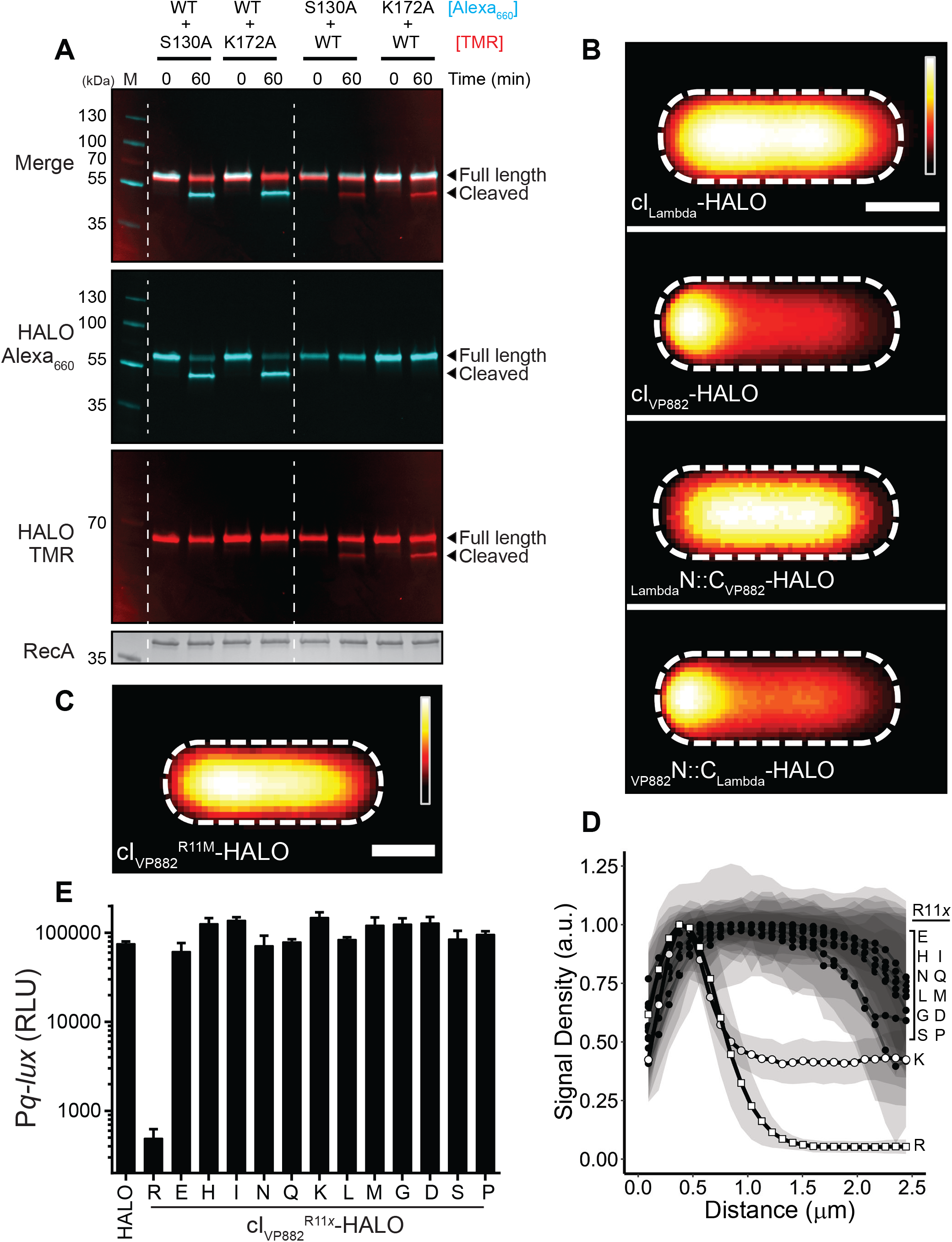
cI_VP882_ is cleaved by an intramolecular mechanism and Qtip recognition occurs at the N-terminus of cI_VP882_. (A) *In vitro* cleavage of WT cI_VP882_-HALO mixed with cI_VP882_ catalytic-site variants monitored by SDS-PAGE analysis and differential labeling. The first lane has the marker, M. The HALO dye combinations used are designated to the right of the proteins. The lanes between the dashed white lines show WT cI_VP882_-HALO conjugated to HALO-Alexa_660_ (cyan) and catalytic-site variants conjugated to HALO-TMR (red). The lanes to the right of the second dashed white line show WT cI_VP882_-HALO conjugated to HALO-TMR (red) and the catalytic-site variants conjugated to HALO-Alexa_660_ (cyan). The gel was imaged under Cy5 and Cy3 filter sets to detect HALO-Alexa_660_ and HALO-TMR, respectively, as designated on the left (see also Figure S1B), prior to being stained for total protein (bottom panel). The composite of the HALO-Alexa_660_ and HALO-TMR channels is shown in the upper-most panel, designated Merge. (B) Composite images from individual cell analyses of *E. coli* producing Qtip and either cI_Lambda_, cI_VP882_, or the indicated chimeras, each fused to HALO and labeled with HALO-TMR. (C) As in (B) with Qtip and cI_VP882_^R11M^-HALO. In (B) and (C), HALO-TMR fluorescence intensity and scale bar represented as in Figure 1D. (D) Line plot of HALO-TMR fluorescence intensity extracted from individual cell images of *E. coli* producing Qtip and either WT cI_VP882_-HALO (designated R) or another cI_VP882_^R11*x*^-HALO variant (designated by the letters on the right). WT cI_VP882_-HALO (open squares) and cI_VP882_^R11K^-HALO (open circles) and the cI_VP882_^R11*x*^-HALO variants (closed circles). Distance along the x-axis as in Figure 1D. Shaded regions represent ± 1 SD from the mean. (E) Light production by *E. coli* harboring the P*q-lux* reporter and a control plasmid (HALO), WT cI_VP882_-HALO (denoted R), or the designated cI_VP882_^R11*x*^-HALO variant. The *x* indicates the amino acid residue at position 11. RLU as in Figure 1. Data represented as mean ± SD with n = 3 biological replicates.

### Qtip Recognizes the DNA-Binding Domain of cI_VP882_ but not that of cI_Lambda_

Our above results show that Qtip prevents cI_VP882_ autoproteolysis, however, Qtip does not depend on the cleavage or catalytic site residues in cI_VP882_ to sequester the protein (see Figure 2D). Moreover, cI_VP882_ has functionally analogous domains to those in cI_Lambda_, and yet, Qtip sequesters cI_VP882_ but not cI_Lambda_ (Silpe and Bassler, 2019a). We thus wondered what determinants are required in a cI protein for recognition by Qtip. We took advantage of the conserved domain architecture between cI_VP882_ and cI_Lambda_ and the similarities in their cleavage mechanisms to engineer two chimeric cI repressors for assessment of interaction with Qtip. Specifically, we exchanged the N-terminal (DNA-binding) and C-terminal (catalytic) domains of cI_VP882_ and cI_Lambda_, leading to _VP882_N∷C_Lambda_-HALO and _Lambda_N∷C_VP882_-HALO proteins (Figure S1A). Both chimeras retained the ability to bind DNA (Figure S2A) and to undergo autoproteolysis (Figure S2B). Moreover, the DNA-binding specificity of each chimera is set by the protein from which the N-terminal DNA-binding portion was derived, i.e., the _VP882_N∷C_Lambda_-HALO chimera, like full length cI_VP882_-HALO, bound phage VP882 DNA but not lambda DNA (Figure S2A). Conversely, the _Lambda_N∷C_VP882_-HALO chimera, while not as stable as the _VP882_N∷C_Lambda_-HALO chimera (see 0’ timepoint in Figure S2B, and Figure S2C), bound lambda DNA, but not VP882 DNA (Figure S2A). We assayed whether Qtip could bind to these chimeras. As controls, and consistent with our earlier findings (Silpe and Bassler, 2019a), we show that the cI_Lambda_-HALO protein was not sequestered by Qtip, whereas localization to the poles occurred for cI_VP882_-HALO (Figure 3B). _Lambda_N∷C_VP882_-HALO, like cI_Lambda_-HALO, remained cytoplasmic in the presence of Qtip, suggesting that Qtip does not interact with cI_Lambda_-HALO or the _Lambda_N∷C_VP882_-HALO chimera. By contrast, the _VP882_N∷C_Lambda_-HALO fusion was sequestered to the poles by Qtip similar to what occurred between Qtip and WT cI_VP882_-HALO (Figure 3B). These results suggest that Qtip recognizes the N-terminal region of cI_VP882_.

### A Subset of the Amino Acid Residues Required for cI_VP882_ to Bind DNA are also Required for Qtip to Bind cI_VP882_

We have shown that Qtip binds the cI_VP882_ N-terminal, DNA-binding domain (Figure 3B) and yet the DNA-binding deficient variant, cI_VP882_^DBD*^-HALO, continues to be sequestered by Qtip (Figure 1D). These results suggest that DNA binding by cI_VP882_ and Qtip binding by cI_VP882_ could be separable functions. To examine this possibility, we sought to identify a cI_VP882_ mutant that was resistant to Qtip-directed localization. We randomly mutagenized WT *cI_VP882_-HALO* on a plasmid and transformed the pool of mutant plasmids into *E. coli* harboring *qtip* under an anhydrotetracyline (aTc)-inducible promoter. When co-produced with Qtip, cI_VP882_^R11M^-HALO remained cytoplasmic, suggesting that mutation of the arginine at position 11 of the cI_VP882_ repressor leads to a loss in Qtip recognition (Figures 3C and S1A). However, a trivial explanation for this phenotype is that the arginine to methionine change at position 11 provides an alternative cI_VP882_-HALO translation start site, and the absence of the first 10 amino acid residues prevents Qtip from binding. To eliminate this possibility, we used semi-arbitrary PCR to engineer random codons at position 11 in cI_VP882_-HALO. In addition to R11M, we recovered cI_VP882_^R11*x*^-HALO [where *x* = D, E, G, H, I, L, N, P, Q, S, K]. All of these alleles remained cytoplasmic in the presence of Qtip except for cI_VP882_^R11K^, which showed partial localization (Figure 3D). These results demonstrate that, in cI_VP882_, the amino acid at position 11 is critical for susceptibility to Qtip. To determine if resistance to Qtip is separable from the DNA-binding function of cI_VP882_, we tested cI_VP882_^R11M^-HALO and the 11 other R11 variants for their ability to repress P*q-lux*. Despite the above cI_VP882_ domain analysis, which did not implicate R11 as being involved in DNA binding (Figure S1A), Figure 3E shows that none of our cI_VP882_^R11*x*^-HALO variants, including cI_VP882_^R11K^-HALO, repressed P*q-lux*. We interpret these data to mean that, at least with respect to amino acid residue R11, elimination of Qtip recognition by cI_VP882_ also eliminates cI_VP882_ DNA-binding activity. Additionally, our findings that cI_VP882_^R11K^-HALO and cI_VP882_^DBD*^-HALO fail to bind DNA but can still be bound by Qtip (Figures 1D and 3D) suggest that Qtip binding to cI_VP882_ requires only a subset of the residues cI_VP882_ uses to bind DNA.

### Qtip Localizes to Cell Poles in the Absence of its Partner Repressor

Having identified the residues on cI_VP882_ that confer its functions and drive its interaction with Qtip, we next sought to pinpoint the amino acids on Qtip that are crucial for its activities. To monitor Qtip localization *in vivo*, we employed a SNAP-Qtip fusion, which enables microscopic visualization and tracking using fluorescent SNAP ligands. We previously showed that SNAP-Qtip produced from an aTc-inducible promoter sequesters cI_VP882_-HALO identically to aTc-induced untagged Qtip (Silpe and Bassler, 2019a). Figures S3A and S3B demonstrate that, also like native Qtip, SNAP-Qtip derepressed cI_VP882_-mediated P*q-lux* expression and induced lysis in VP882 lysogens. These results show that the SNAP tag does not interfere with known Qtip functions and the construct can be used as a fluorescent tool to probe Qtip properties.

The first question we addressed was whether Qtip could transit to the pole in the absence of cI_VP882_ or whether complex formation between Qtip and cI_VP882_ is required. Figure 4A shows that SNAP-Qtip localized to the cell pole irrespective of whether or not cI_VP882_-HALO was present. One possibility is that the Qtip polar localization we observed occurred as a consequence of high-level production from the aTc-inducible promoter. We titrated down the inducer and we could find no concentration of aTc capable of generating measurable SNAP-Qtip signal that did not also cause polar localization (Figure S3C). Natural Qtip levels have not been established so we do not have a comparison. However, since the SNAP tag and cI_VP882_-HALO are both cytoplasmic (Figure 4A and (Silpe and Bassler, 2019a)), our data suggest that polar localization of the Qtip-cI_VP882_ complex is a property conferred by Qtip.

**Figure 4:**
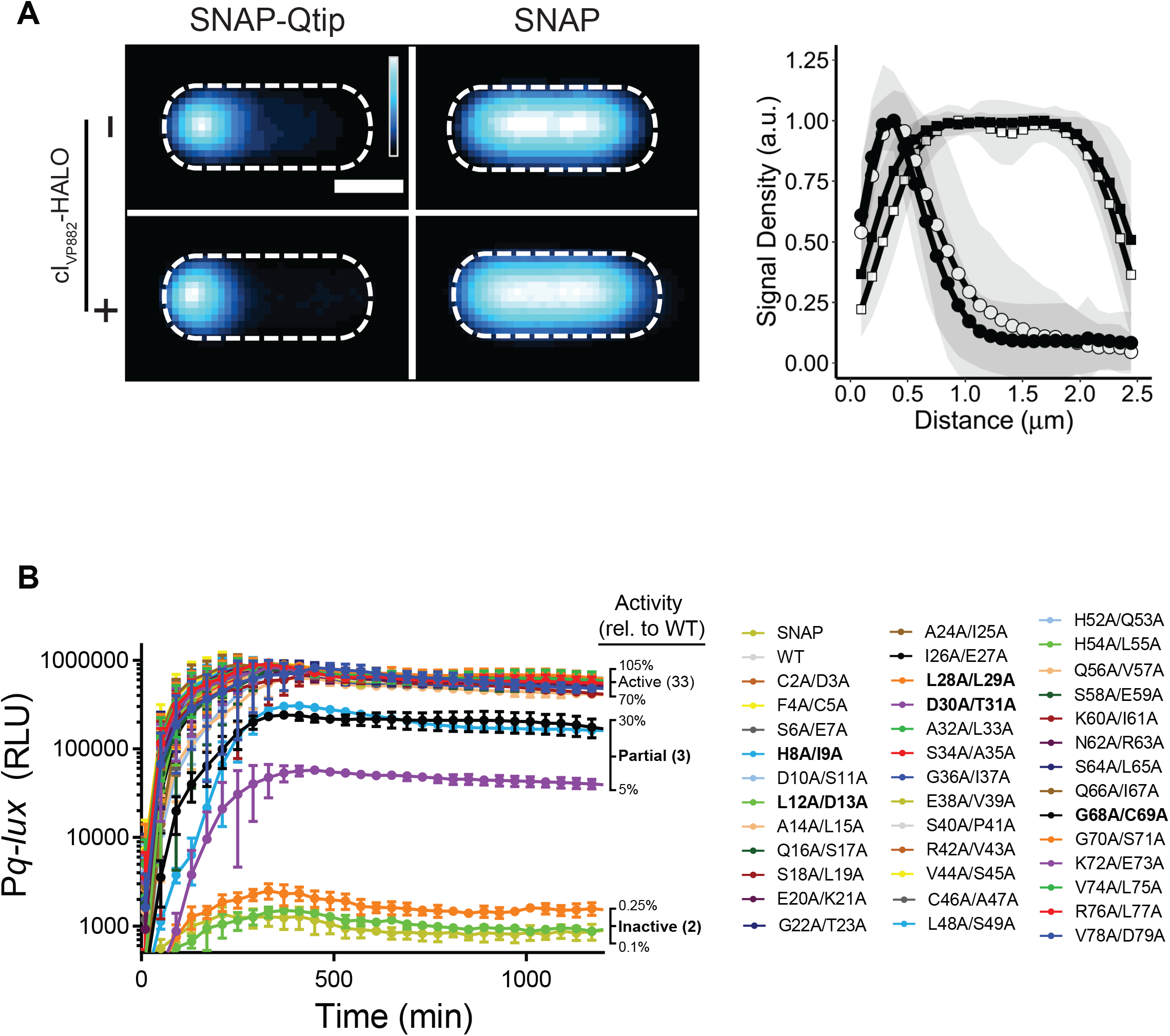
SNAP-Qtip localizes to the poles in the absence of its partner repressor cI_VP882_. (A) Left: Composite images from individual cell analyses of *E. coli* producing SNAP-Qtip or SNAP, in the absence or presence of cI_VP882_-HALO (designated as – or + on the left side of the images) labeled with SNAP-JF_503_. SNAP-JF_503_ fluorescence intensity is displayed as a cyan heat map. Black and white reflect the lowest and highest intensity, respectively. Scale bar, as in Figure 1D. Right: Line plot of SNAP-JF_503_ fluorescence intensity extracted from individual cell images used in the left panel. Distance along the x-axis as in Figure 1D. Symbols: SNAP-Qtip (circles) and SNAP (squares), each in the absence (open) or presence (closed) of cI_VP882_-HALO. Shaded regions represent ± 1 SD from the mean. (B) Time-course of light production from the P*q-lux* reporter in *E. coli* producing cI_VP882_ and either SNAP, WT SNAP-Qtip, or the designated SNAP-Qtip double-alanine variant. Variants exhibiting partial activity or that are inactive are shown in bold text in the key. RLU as in Figure 1. Data represented as mean ± SD with n = 3 biological replicates.

### *Alanine-Scanning Mutagenesis of Qtip Decouples its Localization Function from its Function as an Inhibitor of* cI_VP882_*Repressor Activity*

Our finding that Qtip sequesters and inhibits cI_VP882_ autocleavage and DNA binding led us to wonder whether the ability of Qtip to localize cI_VP882_ is separable from its ability to inactivate cI_VP882_. To explore this issue, we performed alanine scanning mutagenesis on Qtip. We altered two consecutive codons at a time. Qtip is a 79 amino acid protein. Excluding the start and stop codons and one existing Ala-Ala pair, this strategy required generation of 38 Qtip Ala-Ala double mutants, each fused to SNAP and cloned onto a plasmid under the aTc-inducible promoter. We assessed each Qtip allele for inactivation of cI_VP882_ DNA-binding activity and for co-localization with cI_VP882_-HALO.

Regarding cI_VP882_ DNA binding and repressor function: we transformed the plasmids carrying the mutant *SNAP-qtip* alleles into *E. coli* harboring a second plasmid containing *cI_VP882_* and P*q-lux*. The logic was: cI_VP882_ represses P*q*-*lux*, and SNAP-Qtip inactivates cI_VP882_, thus, introduction of a functional SNAP-Qtip mutant will induce light production, while introduction of a non-functional Qtip mutant will not (Figure 1A). Thirty three of the 38 SNAP-Qtip Ala-Ala mutants showed at least ~70% of wild-type (WT) SNAP-Qtip activity in the P*q*-*lux* assay (Figure 4B). Of the 5 remaining SNAP-Qtip mutants, 3 (SNAP-Qtip^H8A/I9A^, SNAP-Qtip^D30A/T31A^, and SNAP-Qtip^G68A/C69A^) exhibited partial activity (5%-30% of WT) and 2 (SNAP-Qtip^L12A/D13A^ and SNAP-Qtip^L28A/L29A^) were incapable of driving light production (<0.3% of WT). Figure S4A shows that 4 of the 5 defective SNAP-Qtip mutant proteins were produced at approximately the same level as WT SNAP-Qtip. Only SNAP-Qtip^L28A/L29A^ was produced at lower than WT levels. Thus, differences in protein production for SNAP-Qtip^H8A/I9A^, SNAP-Qtip^D30A/T31A^, SNAP-Qtip^G68A/C69A^, and SNAP-Qtip^L12A/D13A^ cannot be responsible for their attenuated activity.

Regarding co-localization with cI_VP882_: We used confocal microscopy to assess the localization patterns of the five defective SNAP-Qtip variants and cI_VP882_-HALO by labeling cells in which the proteins had been co-expressed with the red-fluorescent HALO-ligand HALO-TMR and the green fluorescent SNAP-ligand SNAP-JF_503_ (Grimm et al., 2017). Figure S4B shows that the 3 partially-inactive SNAP-Qtip mutants (SNAP-Qtip^H8A/I9A^, SNAP-Qtip^D30A/T31A^, and SNAP-Qtip^G68A/C69A^) were also partially defective in localizing cI_VP882_-HALO to the pole. The 2 SNAP-Qtip mutants that failed to inactivate cI_VP882_-HALO (SNAP-Qtip^L12A/D13A^ and SNAP-Qtip^L28A/L29A^) also failed to localize cI_VP882_-HALO (Figure S5A and B, left panels). Remarkably, however, SNAP-Qtip^L12A/D13A^, while unable to inactivate the cI_VP882_ repressor or to localize cI_VP882_-HALO to the pole, did itself localize to the cell pole (Figure S5A, left panel).

We next investigated whether, in the cases of the 2 Qtip mutants that failed to inactivate cI_VP882_ (L12A/D13A and L28A/L29A), both Ala substitutions or only a single Ala residue were required to confer the mutant phenotype. To do this, we made all of the corresponding single Ala substitutions. In the case of the SNAP-Qtip^L12A/D13A^ variant, the D13A alteration was sufficient for the mutant phenotype: SNAP-Qtip^D13A^ localized to the pole and it did not drive polar localization or inactivation of cI_VP882_ (Figure 5A and 5B). By contrast, SNAP-Qtip^L12A^ exhibited partial polar localization (Figure S5A, right panel) and it had measurable inhibitory activity against cI_VP882_ (Figure 5A).

**Figure 5:**
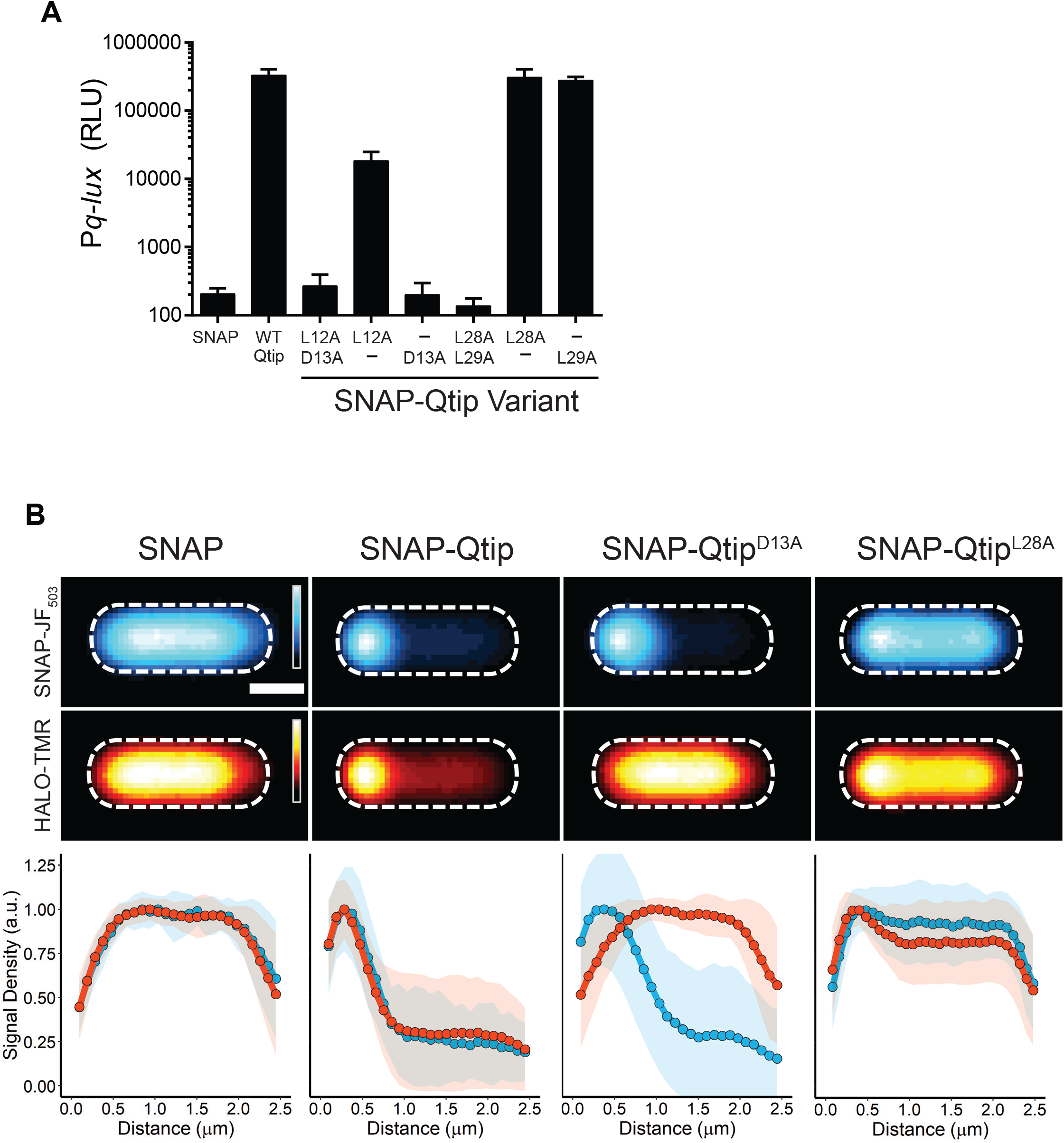
Qtip polar localization and inhibitory activity against cI_VP882_ are separable. (A) Light production from the P*q-lux* reporter in *E. coli* producing cI_VP882_ and SNAP, WT SNAP-Qtip, or the indicated SNAP-Qtip variant. RLU as in Figure 1. Data represented as mean ± SD with n = 3 biological replicates. (B) Upper two rows: Composite images from individual cell analyses of *E. coli* producing cI_VP882_-HALO and either SNAP, SNAP-Qtip, or the indicated SNAP-Qtip variant. Samples labeled with SNAP-JF_503_ (top row) and HALO-TMR (bottom row). Fluorescence intensity displayed as cyan (SNAP-JF_503_) and red (HALO-TMR) heat maps. Scale bar, as in Figure 1D. Lower: Line plots of the SNAP-JF_503_ (blue) and HALO-TMR (red) fluorescence intensities extracted from the single cells. Distance along the x-axis as in Figure 1D. Shaded regions represent ± 1 SD from the mean.

The case of SNAP-Qtip^L28A/L29A^, which was cytoplasmic and inactive against cI_VP882_, was less straightforward (see Figure S5B). The localization defect of SNAP-Qtip^L28A/L29A^ can be explained by mutation of only the first codon: SNAP-Qtip^L28A^ was cytoplasmic (Figure 5B), whereas SNAP-Qtip^L29A^ was partially localized at the pole (Figure S5B). However, the defect in inhibition of cI_VP882_ repressor activity by SNAP-Qtip^L28A/L29A^ cannot be explained by either of the single site variants: both SNAP-Qtip^L28A^ and SNAP-Qtip^L29A^ were active against cI_VP882_-HALO (Figure 5A). In addition, the poor production of SNAP-Qtip^L28A/L29A^ could not be ascribed to either single site variant as both SNAP-Qtip^L28A^ and SNAP-Qtip^L29A^ were produced at the levels of WT SNAP-Qtip (Figure S4A). Our interpretation is that Qtip residue L28 is required for Qtip localization but dispensable for inhibitory activity against cI_VP882_, whereas Qtip residue L29 is dispensable for both traits. In the SNAP-Qtip^L28A/L29A^ double mutant, there is a synthetic defect that causes reduced protein levels, which underpin the SNAP-Qtip^L28A/L29A^ defects against cI_VP882_-HALO. Collectively, our results with the defective Qtip mutants reveal that a single site substitution, D13A, abolishes the ability of Qtip to inactivate cI_VP882_ while not affecting the ability of Qtip to localize to the cell pole, and a single substitution at a different site, L28A, disrupts localization but not inhibitory activity against cI_VP882_.

### The Phage P22 Antirepressor, Ant, Inactivates cI_Lambda_ without Sequestration

Among the first antirepressors discovered was Ant of phage P22 (Botstein et al., 1975; Levine et al., 1975; Susskind and Botstein, 1975). While Ant is approximately four times the size of Qtip (34.6 kDa vs 8.4 kDa) and the proteins have no homology, the P22-Ant system does share similarities with the VP882-Qtip system: In both phages, the repressor protein can both be inactivated by cleavage and can be inactivated by an antirepressor, rather than one or the other. Ant, in addition to inactivating the P22 repressor, called C2, and inducing lysis, also binds to and interferes with DNA binding of cI_Lambda_ (Susskind and Botstein, 1975), suggesting cross-reactivity analogous to what we observe between Qtip and other phage repressors (Silpe and Bassler, 2019a). We thus considered how features of Qtip-mediated repressor inactivation might parallel those of Ant. Specifically, we wondered whether polar localization occurs when Ant inactivates its partner repressors, which to our knowledge, has not yet been investigated.

To verify that Ant affects cI_Lambda_-directed lysis-lysogeny, we cloned the *ant* gene under the aTc-inducible promoter and introduced it into *E. coli* lysogenized by the temperature-sensitive lambda cI857 phage (Sussman and Jacob, 1962). Lambda cI857 only lysogenizes *E. coli* at temperatures at or below 30°C. As a control, and consistent with our above result showing that Qtip does not sequester cI_Lambda_ (see Figure 3B and (Silpe and Bassler, 2019a)), we show that induction of Qtip or SNAP-Qtip production in the lambda cI857 lysogen did not induce lysis, whereas lysis occurred following induction in *E. coli* harboring the VP882 phage as an episome (Figure 6A and 6B). These results demonstrate that Qtip is a functional antirepressor for phage VP882 but not for phage lambda. In contrast, aTc-induced production of Ant inhibited growth of *E. coli* carrying the lambda cI857 lysogen, showing that the cI_Lambda_ protein is inactivated by Ant (Figure 6A). Production of Ant in *E. coli* harboring the VP882 episome did not affect growth (Figure 6B) nor did Ant derepress P*q-lux* expression in the *E. coli* reporter system (Figure S6A). Taken together, these results indicate that Ant is a functional antirepressor for phage lambda but not for phage VP882. Like Qtip, Ant remained functional when fused to SNAP (Ant-3xFLAG-SNAP; Figure 6A and 6B), however, unlike SNAP-Qtip, Ant-3xFLAG-SNAP was cytoplasmic, and its localization did not change in the presence or absence of cI_VP882_-HALO, cI_Lambda_-HALO, or its natural partner, C2-HALO (Figure 6C). Similarly, the presence of Ant-3xFLAG-SNAP did not change the localization of any of the three repressor proteins (cI_VP882_-, cI_Lambda_-, and C2-HALO), and SNAP-Qtip did not change the localization of C2-or cI_Lambda_-HALO (Figure 6C). We conclude that while both Ant and Qtip are capable of inactivating their target and related repressors, Qtip induces polar localization, while Ant does not.

**Figure 6:**
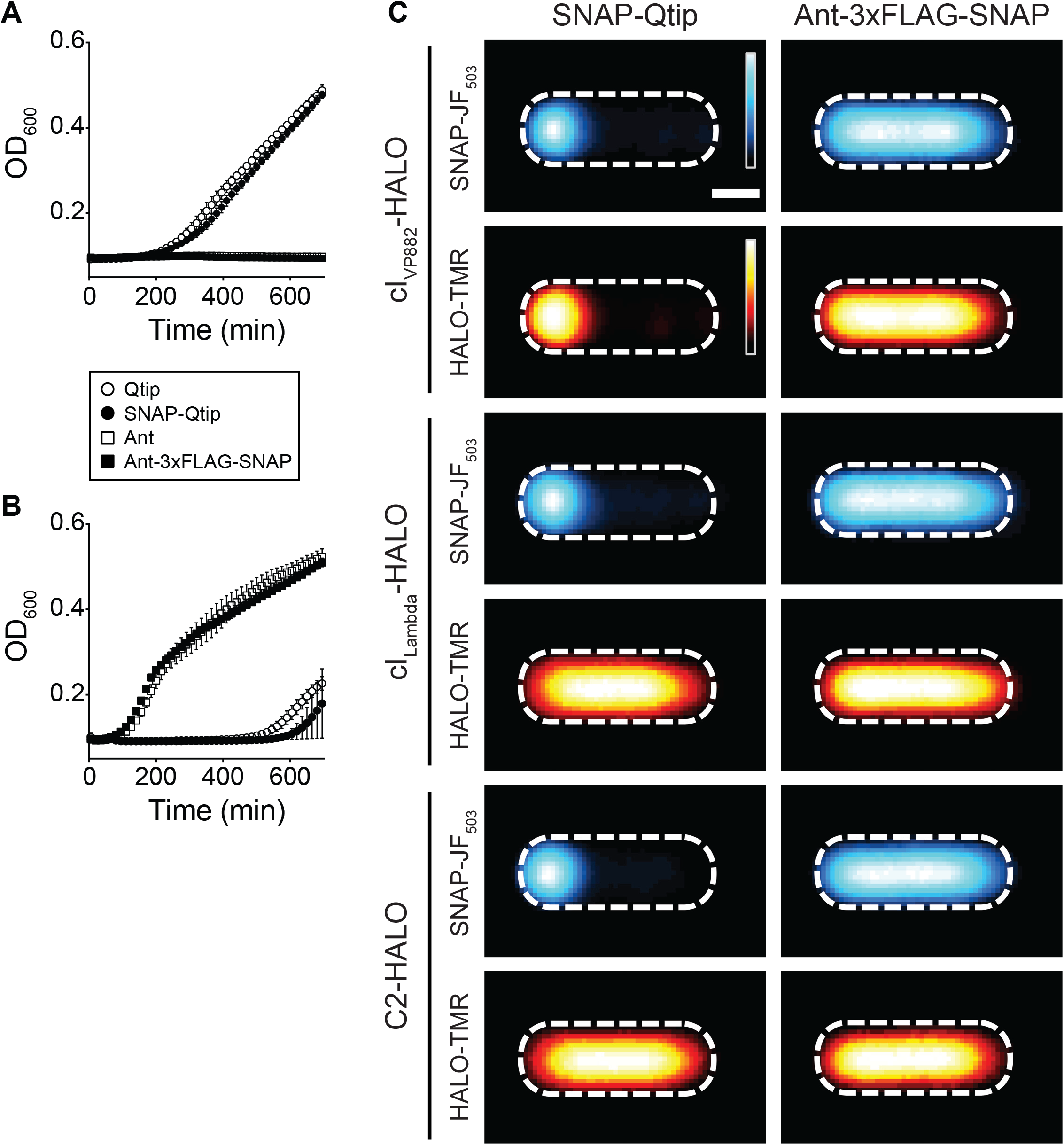
Ant is cytoplasmic and induces lysis in phage lambda but not in phage VP882, and Qtip localizes at the pole and induces lysis in phage VP882 but not in phage lambda. Growth curve of *E. coli* lysogenized by phage lambda cI857 (A) or phage VP882 CmR∷Tn*5* (B), each expressing the antirepressor constructs indicated in the key. (A) was performed at 30°C, (B) at 37°C. Data represented as mean ± SD with n = 3 biological replicates. (C) Composite images from individual cell analyses of *E. coli* producing SNAP-Qtip (left column) or Ant-3xFLAG-SNAP (right column) and cI_VP882_-HALO (first two rows), cI_Lambda_-HALO (third and fourth rows), or C2-HALO (fifth and sixth rows). SNAP and HALO labeled with SNAP-JF_503_ and HALO-TMR, respectively, and displayed using the same heat map color schemes as in Figure 5B. Scale bar, as in Figure 1D.

## Discussion

Our investigation into interactions between Qtip and cI_VP882_ revealed that, in some respects, cI_VP882_ behaves much like cI_Lambda_. Both repressors block lytic development in their respective phages and have SOS-responsive, RecA-activated catalytic domains located at their C-termini that promote intramolecular cleavage reactions. Both *cI_VP882_ and cI_Lambda_* also encode N-terminal DNA binding domains, however, a feature present in cI_VP882_ that enables Qtip binding is absent from cI_Lambda_. Susceptibility to Qtip provides the VP882 phage with a QS-dependent trigger. Specifically, Qtip, which is produced in response to VqmA_Phage_ binding to the host-produced QS AI DPO, prevents cI_VP882_ from binding DNA.

Temperate phages encoding antirepressors typically possess repressor proteins that lack catalytic C-terminal domains, presumably because the antirepressor fulfills that function. Thus, repressors in these systems are often half the size of cI_Lambda_ (Lemire et al., 2011). cI_VP882_ and P22 C2 are curious in this regard because both possess autoproteolytic activity and are subject to inactivation by antirepressors (Qtip and Ant, respectively). Strikingly, early work on phage P22 showed that an Ant-overproducing P22 mutant that was exposed to UV light to induce RecA-dependent autoproteolysis did not undergo autoproteolysis, which is unlike a similarly treated non-Ant-overproducing P22 prophage (Prell and Harvey, 1983). This experiment suggested that Ant production “protects” C2 from autoproteolysis (Prell and Harvey, 1983). The experiment did not address whether the effect was direct or indirect. Our result showing that Qtip binding inhibits cI_VP882_ autoproteolysis *in vitro* is consistent with the Ant-C2 system, and furthermore demonstrates that antirepressors can directly alter the proteolytic susceptibility of the proteins they bind. We wonder whether antirepressor-directed inhibition of proteolytic degradation of phage repressors has implications for phage-host biology. While untested, we imagine that if antirepressor binding blocks access of RecA* to the repressor, there could be a larger pool of RecA* available to function in other crucial host SOS activities during stress (e.g., LexA cleavage and homologous DNA recombination). In support of this notion, production of the C-terminal domain of cI_Lambda_ inhibits SOS induction of the LexA-controlled regulon, suggesting the phage repressor titrates out the available RecA*, prohibiting it from performing its other functions (Ghodke et al., 2019).

In addition to inhibition of cI_VP882_, Qtip localizes cI_VP882_ to the cell poles. We show that inhibition and polar localization can be separated by single site mutations in Qtip that allow inhibition without localization (L28A) and vice-versa (D13A). Our objective in the mutant screen was to identify sites on Qtip that were absolutely required for Qtip function. Thus, we induced *SNAP-qtip* expression with a saturating amount of aTc to drive high level SNAP-Qtip production. Excluding the three double mutants with intermediate phenotypes (Figures 4B and S4B), this strategy revealed SNAP-Qtip^L12A/D13A^, SNAP-Qtip^D13A^, and SNAP-Qtip^L28A/L29A^. Other SNAP-Qtip point mutants might exhibit interesting but more subtle defects if we performed the experiment using sub-maximal levels of inducer to lower the concentration of SNAP-Qtip produced.

We also consider how the properties we discovered for Qtip compare to other antirepressors. Despite our data showing that the prototypical antirepressor, Ant, does not exhibit polar localization, to our knowledge, only one other phage antirepressor has been investigated at the single cell level and it, like Qtip, does localize to the poles (Davis et al., 2002). Specifically, the RS1 satellite phage to CTXφ of *V. cholerae* encodes the antirepressor, RstC, that binds to the CTXφ prophage repressor, RstR, rendering RstR insoluble. The consequence is production of cholera toxin and the transmission of the satellite and CTXφ phages to new cells (Davis et al., 2002). While RstC and Qtip have no homology, RstC is also a small protein (8.3 kDa) and it also induces aggregation of RstR (Davis et al., 2002). Unlike cI_VP882_, however, RstR does not have an obvious catalytic domain or cleavage site. Also similar to the phage VP882 system, CTXφ is induced by DNA damage via the host SOS response. However, unlike phage VP882, in CTXφ, regulation occurs via LexA binding to phage DNA to repress the expression of lytic genes (Quinones et al., 2005). Derepression occurs upon cleavage of LexA.

The biological significance of localization of Qtip and other antirepressors and the Qtip-directed sequestration of the repressor are not yet clear with respect to the host or the phage. One possibility is that localization ensures a high concentration of Qtip at the poles, which is where phage attachment and injection often occur. Specifically, tracking of quantum dot labeled bacteriophages, including one infecting *V. cholerae*, showed that phage adsorption occurs predominantly at the poles of the host cell (Edgar et al., 2008). Moreover, the FtsH protein of *E. coli*, which is involved in the regulation of lambda cII and cIII stability, is also localized to the poles (Edgar et al., 2008). One model proposed is that localization of FtsH at the site where lambda DNA first appears positions FtsH to regulate lysis-lysogeny of newly injected phage genomes (Edgar et al., 2008). Our work, showing that Qtip is also localized to the poles could implicate it in the superinfection process: the high local concentration of Qtip at the poles could enable it to rapidly act on repressors produced by newly infecting phages. While our current work shows that Ant of P22 does not share the same localization behavior as Qtip, it is worth noting that Ant is thought to be expressed early after infection, and the notion is that Ant plays a role during superinfection (Wu et al., 1987).

*V. cholerae* and *Pseudomonas aeruginosa* strains harboring QS-null mutations are frequently isolated (Feltner et al., 2016; Hammer and Bassler, 2003; Stutzmann and Blokesch, 2016) suggesting that there is pressure to avoid producing or to cheat-on QS-produced public goods. As phage VP882 has the capacity to activate lysis in response to host DPO production, we wondered whether mutations in the phage-encoded QS pathway similarly arise. As one example, we identified a curious 8.5 kb contig (GenBank: NNHH01000051.1) in a recent dataset from mixed *V. parahaemolyticus* populations (Yang et al., 2019) (Figure S6B). The contig spans the region where *vqmA_Phage_* and *qtip* are located in phage VP882, between two conserved genes, *repA* and *telN* (Silpe and Bassler, 2019a). Aligned across its entire length, the contig is largely identical to VP882: the contig-derived RepA and TelN products have 99.2% (1229/1239) and 99.3% (534/538) amino acid identity, respectively, to the proteins encoded in phage VP882 (Figure S6B). However, between *repA* and *telN*, there is an 819 bp deletion on the contig that eliminates *vqmA_Phage_*, *qtip*, and the promoters and ribosome binding sites for both ORFs, leaving only the DNA encoding the first 36 amino acids of Qtip and the first 201 amino acids of VqmA_Phage_ (Figure S6B). If this contig corresponds to a functional VP882-like phage, it suggests that the phage would be unresponsive to DPO and cannot execute QS-induced lysis. One can imagine scenarios in which inactivation of phage QS could benefit the host and/or the phage. For example, lack of the ability to induce host cell-density-dependent lysis could promote longer-term lysogeny, as has been shown to be prevalent for phages in gut ecosystems (Kim and Bae, 2018).

It is also worth noting that phage VP882 is a plasmid-like phage, which we previously showed allows it to be transformed and maintained in bacteria that likely fall outside of the natural host range of the phage (e.g., *Salmonella typhimurium* and *E. coli*) (Silpe and Bassler, 2019a). Perhaps existing as a plasmid enables multiple mechanisms of transfer to new cells. We suspect that bacterial species within the natural phage VP882 host range can contract VP882 as a phage, via infection, or they can be transformed, while species outside of the natural host range can take-up VP882 as a plasmid, only via transformation. Interestingly, a recently deposited NCBI entry (GenBank: AAEKWQ010000030.1) of *Salmonella paratyphi B* variant L(+) tartrate(+) isolated from a human stool sample (CDC, National *Salmonella* Reference Laboratory) revealed a ~37.5 kb contig with ~90% identity (35,422 of 37,472 sites) on a nucleotide basis across its entire length to the genome of phage VP882 (Figure S6C). The *Salmonella*-derived contig has intact *vqmA_Phage_* and *qtip* genes, each with 99.1% (233/235 sites) and 100% (79/79 site) identity at the protein level, respectively, to those of phage VP882. *Vibrios* encode the DPO receptor and effector, VqmA and VqmR, respectively, and, as far as we are aware, *Salmonella* do not. Members from both bacterial families produce DPO (Papenfort et al., 2017). It is therefore possible that phage VP882, by encoding a receptor to a widely produced AI, can use the information encoded in DPO even in bacteria that, themselves, cannot.

In terms of the origin of the *Salmonella*-derived VP882-like contig, we imagine that since one area in which *Salmonella* and *Vibrios* come into contact is in the human host, it is plausible that the phage is transferred between species as a plasmid during human infection by the bacteria. In support of such a model, recent work showed that within the human gut, *Salmonella* persister cells promote the spread of broad-range resistance plasmids even in the absence of their selection (Bakkeren et al., 2019).

It is reported that as many as 80% of phage-encoded gene products are unrelated to proteins of known functions (Hatfull and Hendrix, 2011). Discovering the functions of uncharacterized phage products reveals the rich diversity of processes that phages can affect, including bacterial physiology, phage-phage interactions, and, in phages carried by endosymbiotic bacteria, the eukaryotic host (Perlmutter et al., 2019). Our results provide insight into one mechanism by which a newly discovered phage-encoded protein, Qtip, can function to control phage-bacterial interactions. More generally, based on our findings with phage VP882, we propose that plasmid-like phages have greater host flexibility than integrating phages, and we speculate that their transfer may occur between pathogens in the human host setting.

## Supplemental Information

Supplemental information includes 4 tables and 6 figures and can be found with this article online.

## Acknowledgments

This work was supported by the Howard Hughes Medical Institute, NIH Grant R37GM065859, National Science Foundation Grant MCB-1713731 (to B.L.B.), NIGMS grant T32GM007388 (O.P.D), the Damon Runyon Fellowship Award, DRG-2302-17 (to A.A.B), a Charlotte Elizabeth Procter Fellowship provided by Princeton University (to J.E.S.), and a National Defense Science and Engineering Graduate Fellowship supported by the Department of Defense (to J.E.S). We are grateful to Dr. Luke Lavis for providing ample SNAP-JF_503_ ligand. We thank members of PulseNet and the National *Salmonella* Reference Laboratory for information on the reported *Salmonella*-derived contig. We thank Tom Silhavy for helpful discussions and Betsy Hart for the cI857 strain.

## Author Contributions

J.E.S., D.R.C., and O.P.D. constructed strains; J.E.S., A.A.B., X.H., D.R.C., and O.P.D. performed experiments; J.E.S., A.A.B., X.H., O.P.D., and B.L.B. analyzed data; J.E.S., A.A.B. and B.L.B. designed experiments; J.E.S and B.L.B. wrote the paper.

