## Supplementary material for "Separating Functions of the Phage-Encoded Quorum-Sensing-Activated Antirepressor Qtip": SI Figures and Methods

### FIGURE S1

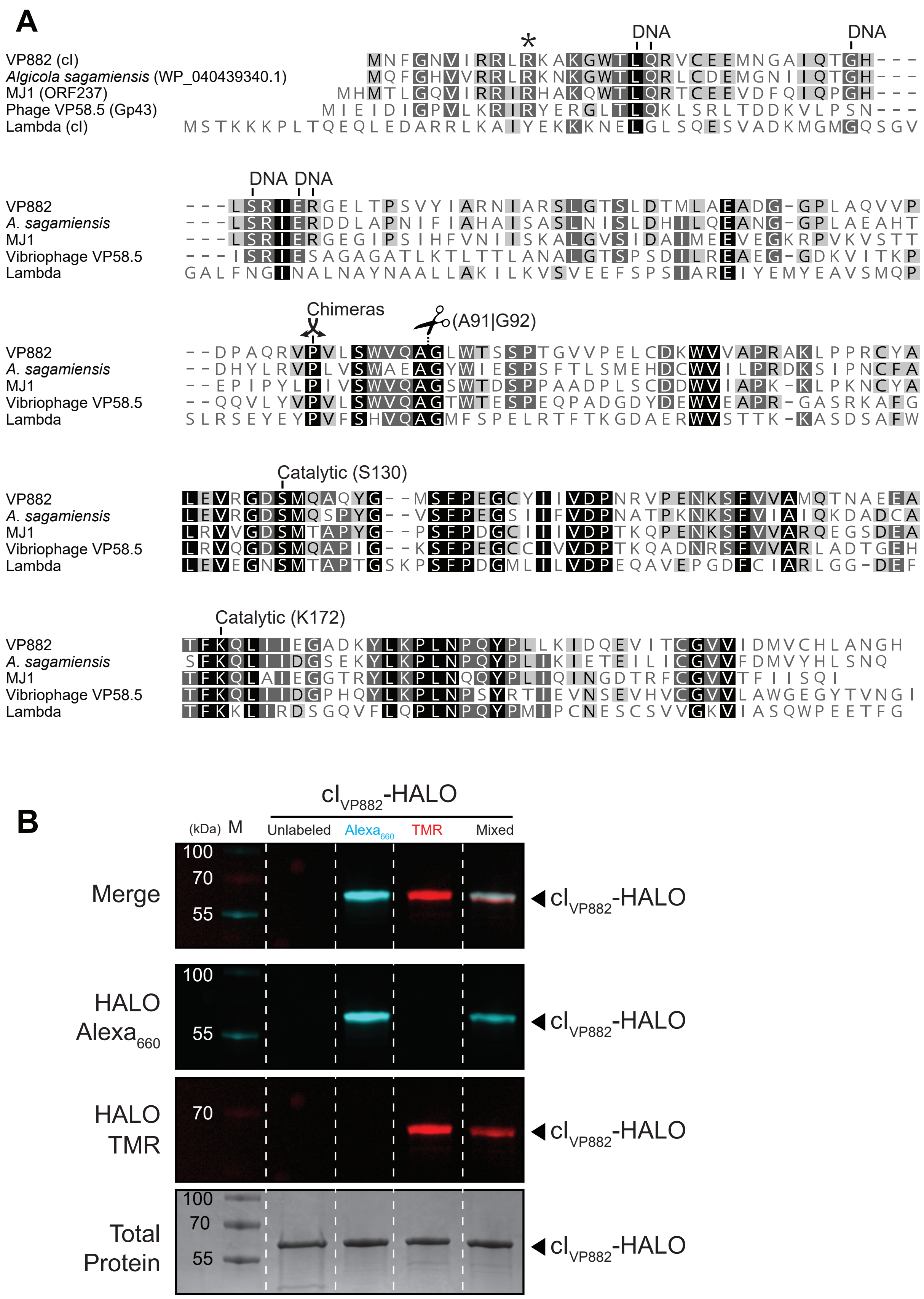

A
*
DNA
DNA
VP882 (cI) MNFGNVIRRLRKAKGWTLQRVCEEMNGAIQTGH---
Algicola sagamiensis (WP_040439340.1) MQFGHVVRRLRKNKGWTLQRLCDEMGNIIQTGH---
MJ1 (ORF237) MHMTLGQVIRRIRHAKQWTLQRTCEEVDFQIQPGH---
Phage VP58.5 (Gp43) MIEIDIGPVLKRIRYERGLTLQKLSRLTDDKVLPSN---
Lambda (cI) MSTKKKPLTQEQLEDARRLKAIYEKKKNELGLSQESVADKMGMGQSGV
DNA DNA
VP882 ---LSRIERGELTPSVYIARNIARSLGTSLDTMLAEADG-GPLAQVVP
A. sagamiensis ---LSRIERDDLAPNIFIAHAISASLNISLDHILQEANG-GPLAEAHT
MJ1 ---LSRIERGEGIPSIHFVNIISKALGVSIDAIMEEVEGKRPVKVSTT
Vibriophage VP58.5 ---ISRIESAGAGATLKTLTTLANALGTSPSDILREAEG-GDKVITKP
Lambda GALFNGINALNAYNAALLAKILKVSVEEFSPSIAREIYEMYEAVSMQP
Chimeras
(A91|G92)
VP882 --DPAQRVPVLSWVQAGLWTSSPTGVVPELCDKWVVAPRAKLPPRCYA
A. sagamiensis --DHYLRVPLVSWAEAGYWIESPSFTLSMEHDCWVILPRDKSIPNCFA
MJ1 --EPIPYLPIVSWVQAGSWTDSPPAADPLSCDDWVIAPK-KLPKNCYA
Vibriophage VP58.5 --QQVLYVPVLSWVQAGTWTESPEQPADGDYDEWVEAPR-GASRKAFG
Lambda SLRSEYEYPVFSHVQAGMFSPELRTFTKGDAERWVSTTK-KASDSAFW
Catalytic (S130)
VP882 LEVRGDSMQAQYG--MSFPEGCYIIVDPNRVPENKSFVVAMQTNAEEA
A. sagamiensis LEVRGDSMQSPYG--VSFPEGSIIFVDPNATPKNKSFVIAIQKDADCA
MJ1 LRVVGDSMTAPYG--PSFPDGCIIIVDPTKQPENKSFVVARQEGSDEA
Vibriophage VP58.5 LRVQGDSMQAPIG--KSFPEGCCIVVDPTKQADNRSFVVARLADTGEH
Lambda LEVEGNSMTAPTGSKPSFPDGMLILVDPEQAVEPGDFCIARLGG-DEF
Catalytic (K172)
VP882 TFKQLIIEGADKYLKPLNPQYPLLKIDQEVITCGVVIDMVCHLANGH
A. sagamiensis SFKQLIIDGSEKYLKPLNPQYPLIKIETEILICGVVFDMVYHLSNQ
MJ1 TFKQLAIEGGTRYLKPLNQQYPLIQINGDTRFCGVVTFIISQI
Vibriophage VP58.5 TFKQLIIDGPHQYLKPLNPSYRTIEVNSEVHVCGVVLAWGEGYTVNGI
Lambda TFKKLIRDSGQVFLQPLNPQYPMIPCNESCSVVGKVIASQWPEETFG
B
cI VP882-HALO
(kDa) M
Unlabeled
Alexa 660
TMR
Mixed
100
70
55
Merge
cI VP882-HALO
HALO Alexa 660
cI VP882-HALO
HALO TMR
cI VP882-HALO
Total Protein
cI VP882-HALO

**Figure S1:** Sequence alignment of $cI_{VP882}$ and other cI-type repressors, and differential HALO-tag labeling to detect proteins by SDS-PAGE analysis, Related to Figures 1, 2, and 3.

(A) Multiple sequence alignment of $cI_{VP882}$ with three related cI-type repressors and $cI_{Lambda}$. Residues in $cI_{VP882}$ predicted to participate in DNA binding are designated "DNA"; putative catalytic-sites are designated "Catalytic", the cleavage site, between a conserved alanine and glycine residue, is represented with scissors (see Figure 2C), sites in $cI_{VP882}$ and $cI_{Lambda}$ used to fuse domains for chimera construction are represented with crossed-arrows (see Figure 3B), and a site required for Qtip recognition is represented with an asterisk (see Figure 3C). Box shading indicates residues that are identical across all (black), the same in 4 of 5 (dark gray), or in 3 of 5 (light gray) proteins. (B) SDS-PAGE analysis of unlabeled $cI_{VP882}$-HALO, $cI_{VP882}$-HALO conjugated to HALO-$Alexa_{660}$ (cyan), $cI_{VP882}$-HALO conjugated to HALO-TMR (red), or when the separately labeled proteins were combined. $Alexa_{660}$ but not TMR-labeled proteins are detected with the Cy5 filter set, designated HALO-$Alexa_{660}$. TMR- but not $Alexa_{660}$- labeled proteins are detected with the Cy3 filter set, designated HALO-TMR. The composite of the HALO-$Alexa_{660}$ and HALO-TMR channels is shown in the upper-most panel, designated Merge. Unlabeled $cI_{VP882}$-HALO is not detected using Cy3 or Cy5 filter sets. All proteins can be visualized by staining the gel for total protein with Coomassie Brilliant Blue, designated Total Protein. Molecular weight marker is designated M.

**FIGURE S2**

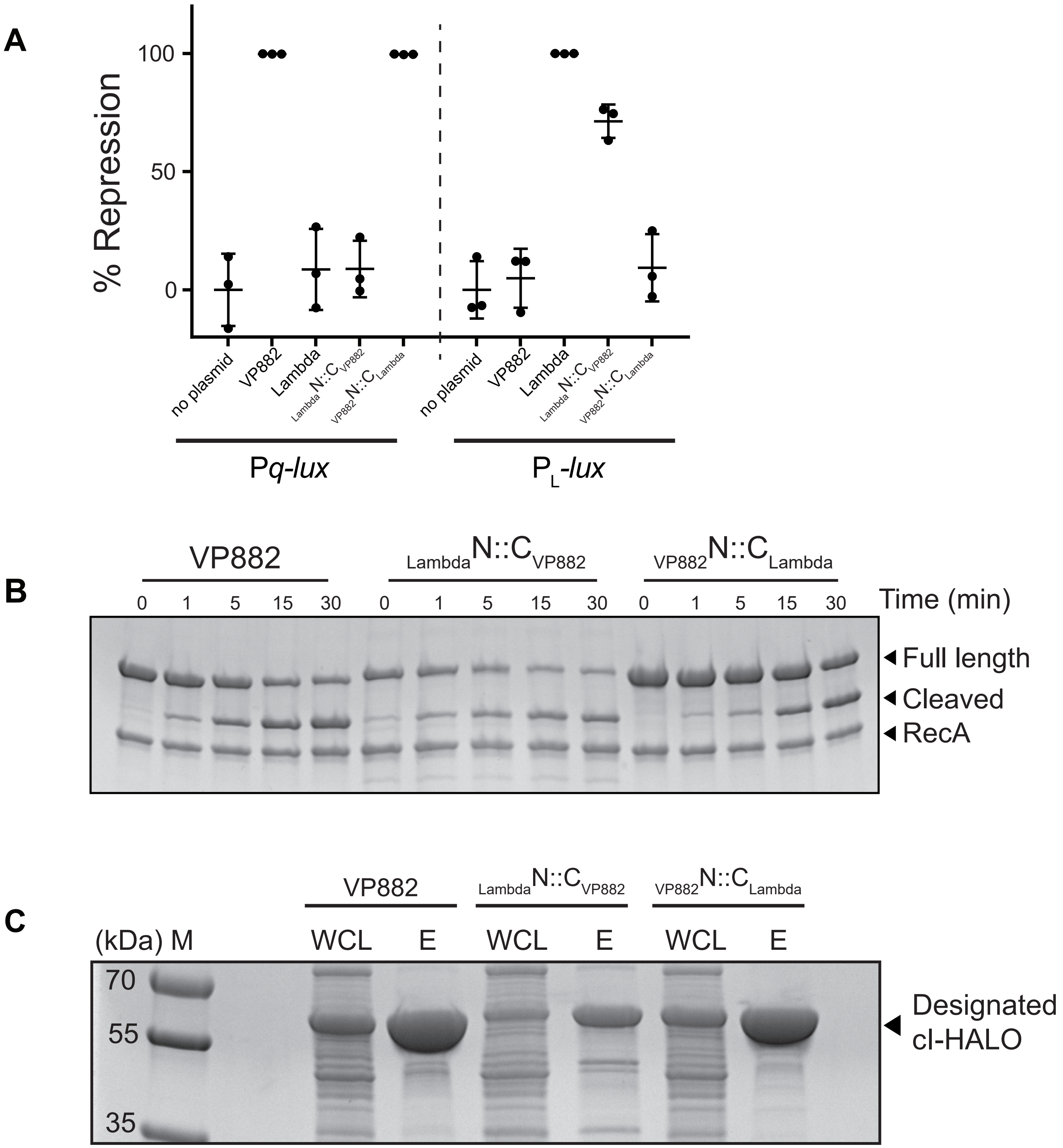

A
% Repression
100
50
0
no plasmid
VP882
Lambda
LambdaN::CVP882
VP882N::CLambda
Pq-lux
PL-lux
B
VP882
LambdaN::CVP882
VP882N::CLambda
0 1 5 15 30
Time (min)
Full length
Cleaved
RecA
C
(kDa) M
WCL
E
70
55
35
Designated
cI-HALO

**Figure S2:** Chimeric cI repressors bind DNA and undergo cleavage *in vitro*, Related to Figure 3

(A) Repression of reporter expression by full-length and chimeric cI repressors. Left side, percent repression of the VP882-encoded P*q*-*lux* reporter or, right side, lambda-encoded $P_L$-*lux* reporter in *E. coli* lacking (no plasmid) or containing the designated cI repressor. %Repression is the difference in RLU obtained for each construct compared to the no-plasmid control. Data represented as mean ± SD with n = 3 biological replicates. (B) *In vitro* cleavage of $cI_{VP882}$-HALO and the chimeric fusions conjugated to HALO-TMR, monitored by SDS-PAGE analysis. Incubation times are noted above each lane. (C) SDS-PAGE analysis of whole-cell lysates (designated WCL) or purified proteins (designated E for eluate) used in (B). Note the lower yield and the presence of possible cleavage products of $_{Lambda}N{::}C_{VP882}$-HALO indicating possible protein instability. Molecular weight marker is designated M.

#### FIGURE S3

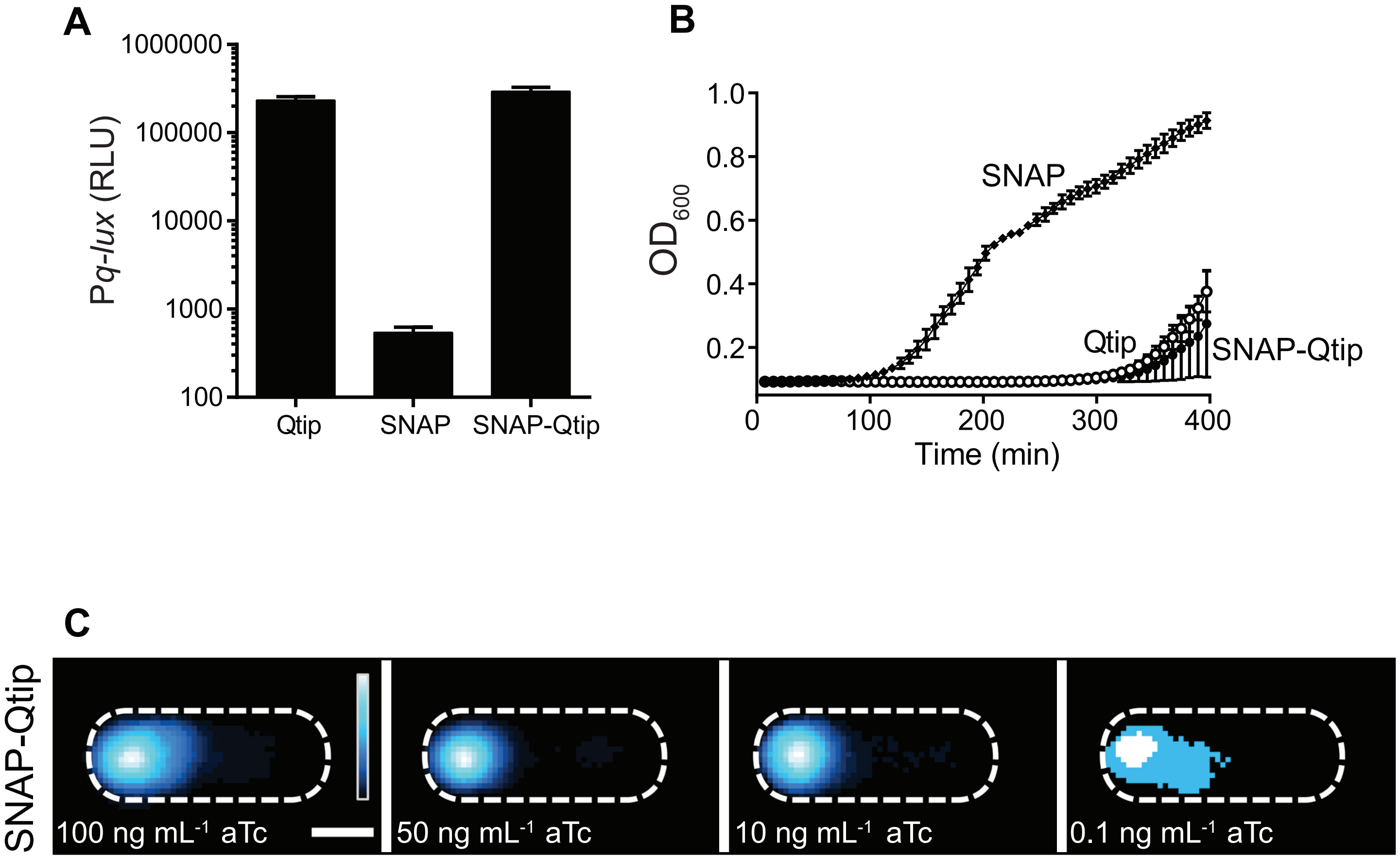
A
1000000
100000
10000
1000
100
Pq-lux (RLU)
Qtip
SNAP
SNAP-Qtip
B
1.0
0.8
0.6
0.4
0.2
OD600
SNAP
Qtip
SNAP-Qtip
0
100
200
300
400
Time (min)
C
SNAP-Qtip
100 ng mL-1 aTc
50 ng mL-1 aTc
10 ng mL-1 aTc
0.1 ng mL-1 aTc

**Figure S3:** The SNAP tag does not interfere with Qtip function and SNAP-Qtip localizes at the cell poles independent of the concentration of inducer, Related to Figure 4.

(A) Light production from the P*q-lux* reporter in *E. coli* producing $cI_{VP882}$ and WT Qtip, SNAP, or SNAP-Qtip. (B) Growth curve of a *Vibrio parahaemolyticus* VP882 lysogen producing the same constructs from (A); SNAP (diamonds), Qtip (open circles), and SNAP-Qtip (closed circles). Data in (A) and (B) represented as mean ± SD with n = 3 biological replicates. (C) Composite images from individual cell analyses of *E. coli* harboring aTc-inducible SNAP-Qtip induced with the indicated amount of aTc. Samples labeled with SNAP-$JF_{503}$ and displayed as in Figure 4A. As noted in the Methods, all images are internally contrasted. The pixelated appearance of the SNAP in the panel showing the 0.1 ng $mL^{-1}$ aTc concentration is a consequence of low induction of SNAP fluorescence. Scale bar, as in Figure 1D.

### FIGURE S4

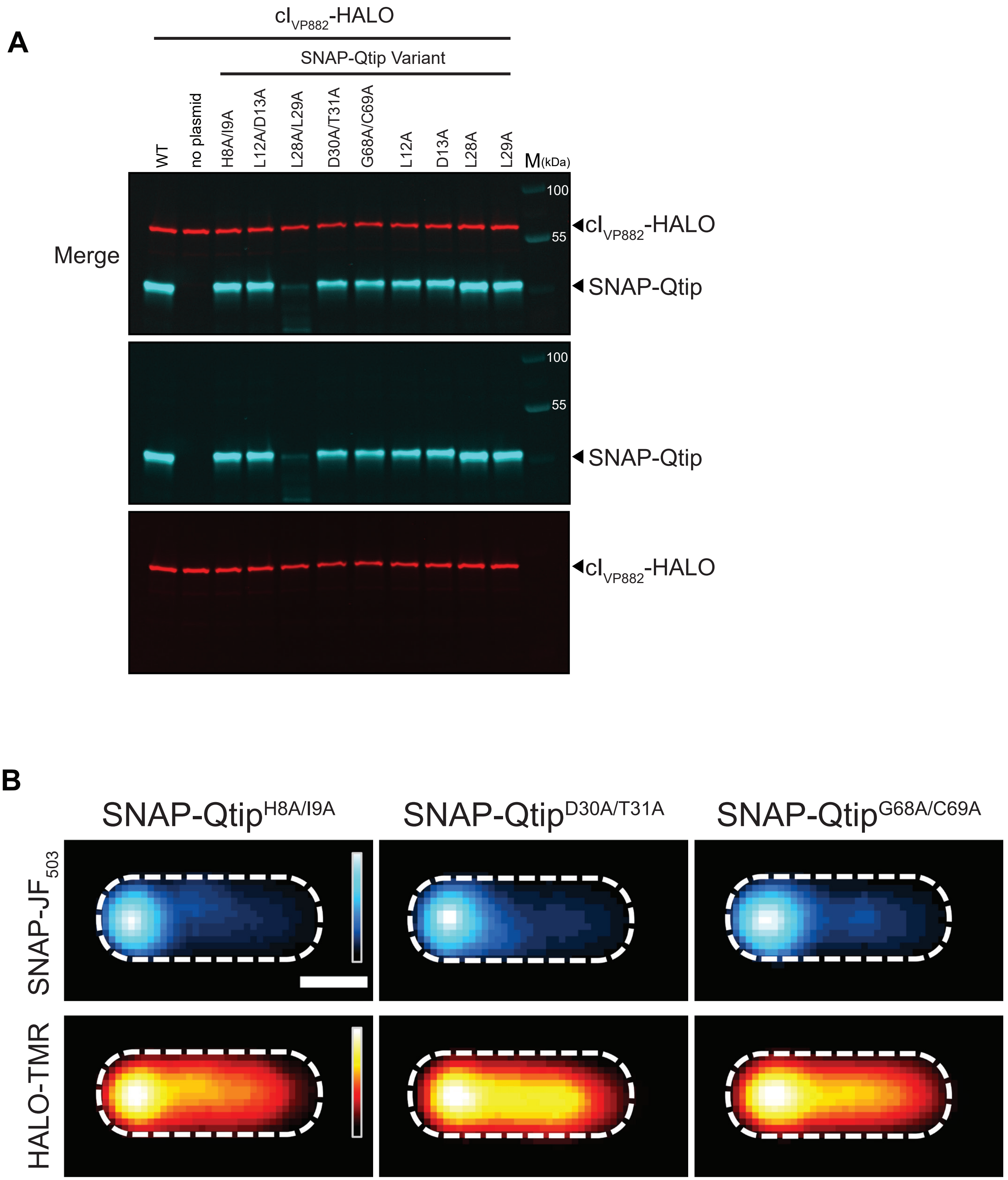
A
cI$_{VP882}$-HALO
SNAP-Qtip Variant
WT
no plasmid
H8A/I9A
L12A/D13A
L28A/L29A
D30A/T31A
G68A/C69A
L12A
D13A
L28A
L29A
M(kDa)
100
55
Merge
cI$_{VP882}$-HALO
SNAP-Qtip
B
SNAP-Qtip$^{H8A/I9A}$
SNAP-Qtip$^{D30A/T31A}$
SNAP-Qtip$^{G68A/C69A}$
SNAP-JF$_{503}$
HALO-TMR

**Figure S4:** Production of WT SNAP-Qtip and SNAP-Qtip variants, and microscopic analysis of SNAP-Qtip$^{H8A/I9A}$, SNAP-Qtip$^{D30A/T31A}$, and SNAP-Qtip$^{G68A/C69A}$, Related to Figures 4 and 5.

(A) Multi-label SDS-PAGE analysis of *E. coli* harboring one plasmid encoding cI$_{VP882}$-HALO and either a second plasmid carrying WT SNAP-Qtip (leftmost lane), no plasmid (second lane) or the indicated SNAP-Qtip variant. SNAP-Qtip was labeled with the far-red fluorescent dye SNAP-Cell 647-SiR and cI$_{VP882}$-HALO was labeled with HALO-TMR. SNAP-Cell 647-SiR (cyan channel) and HALO-TMR (red channel) signals were visualized by imaging the gel under the Cy5 and Cy3 filter sets, respectively, followed by overlaying the channels into a single image (Merge). Molecular weight marker is designated M. (B) Composite images from individual cell analysis of *E. coli* producing cI$_{VP882}$-HALO and either SNAP-Qtip$^{H8A/I9A}$, SNAP-Qtip$^{D30A/T31A}$, or SNAP-Qtip$^{G68A/C69A}$. Samples labeled with SNAP-JF$_{503}$ and HALO-TMR, and displayed as in Figure 5B. Scale bar, as in Figure 1D.

**FIGURE S5**

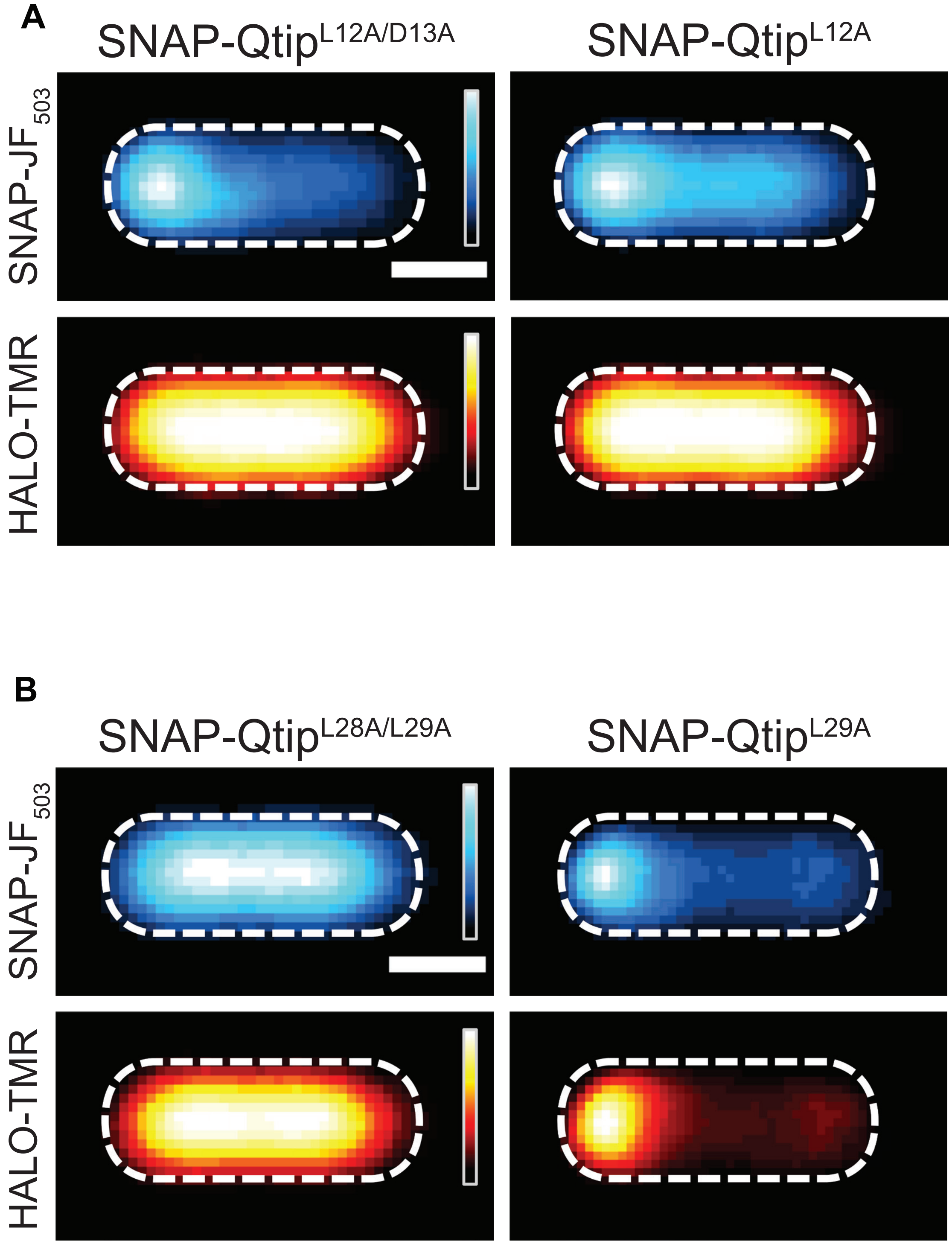
A
SNAP-Qtip$^{L12A/D13A}$
SNAP-Qtip$^{L12A}$
SNAP-JF$_{503}$
HALO-TMR
B
SNAP-Qtip$^{L28A/L29A}$
SNAP-Qtip$^{L29A}$
SNAP-JF$_{503}$
HALO-TMR

**Figure S5:** Microscopic analysis of SNAP-Qtip$^{L12A/D13A}$, SNAP-Qtip$^{L12A}$, SNAP-Qtip$^{L28A/L29A}$, SNAP-Qtip$^{L29A}$, Related to Figures 4 and 5.

Composite images from individual cell analyses of *E. coli* producing $cI_{VP882}$-HALO and either SNAP-Qtip$^{L12A/D13A}$, SNAP-Qtip$^{L12A}$ (A), or SNAP-Qtip$^{L28A/L29A}$, SNAP-Qtip$^{L29A}$ (B). Samples labeled with SNAP-$JF_{503}$ and HALO-TMR, and displayed as in Figure 5B. Scale bars, as in Figure 1D.

#### FIGURE S6

A

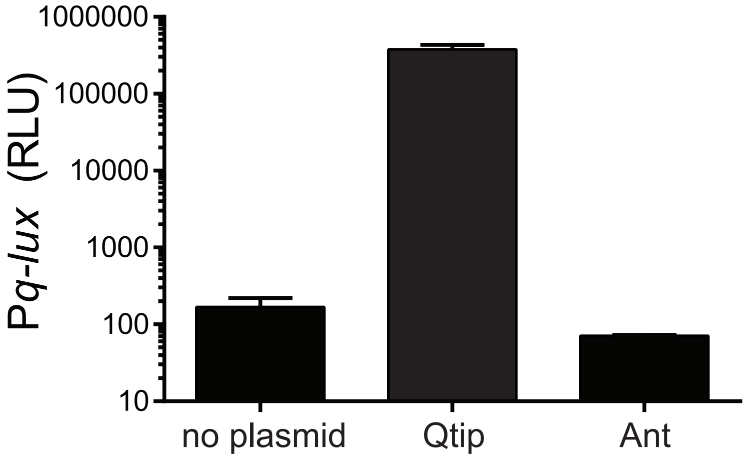


B

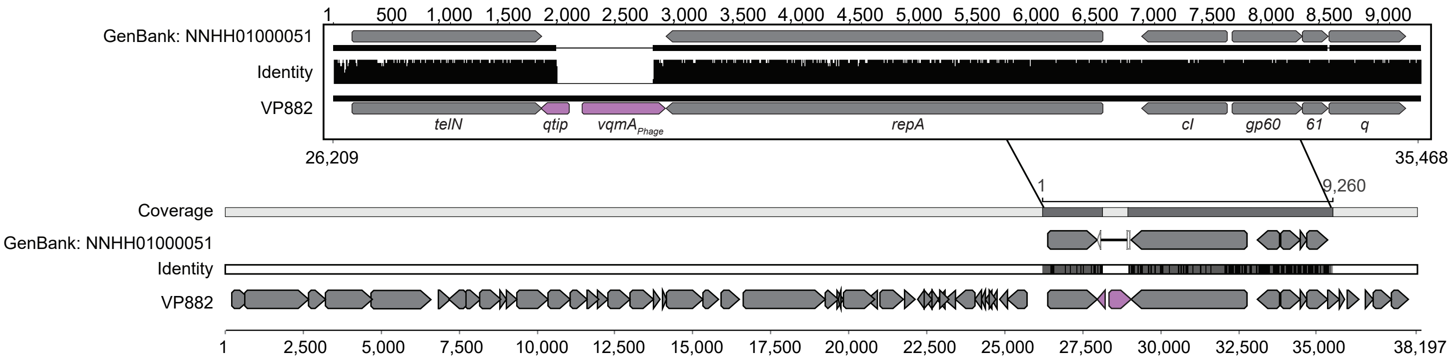


C

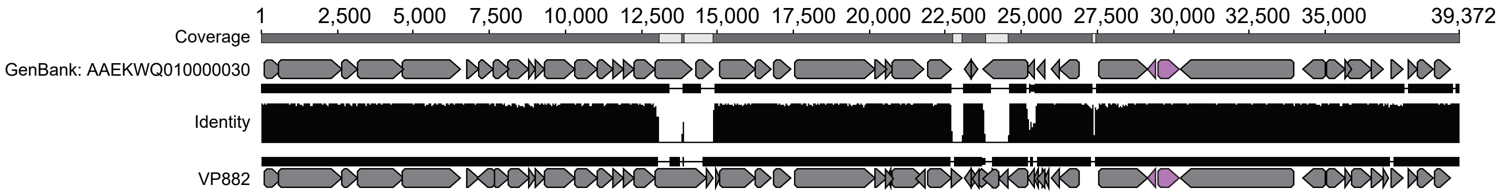

**Figure S6:** Ant does not induce P*q-lux* expression, and phage VP882-like elements are present in *Vibrio* and *Salmonella* isolates, Related to Figure 6.

(A) Light production from the P*q-lux* reporter in *E. coli* producing $cI_{VP882}$ (no plasmid) or $cI_{VP882}$ and a plasmid encoding Qtip or Ant. RLU as in Figure 1. Data represented as mean ± SD with n = 3 biological replicates. (B) Alignment of GenBank: NNHH01000051 across the VP882 phage genome. Arrows indicate ORFs, purple indicates *qtip* and $vqmA_{Phage}$. Top inset, magnified region of homology across the region containing *qtip* and $vqmA_{Phage}$. Identity plotted between the two alignments with a sliding window size of 1. The gap in the inset indicates the 819 bp deletion in the NNHH01000051 contig. Numbering below and above the alignment indicate the genomic coordinates in phage VP882 and the consensus sequence in the contig, respectively. (C) Alignment of the *Salmonella*-derived element (GenBank: AAEKWQ010000030) across the phage VP882 phage genome. Numbering above the alignment indicates coordinates in the consensus sequence. Identity, gaps, and color scheme as in (B).

#### Method Details

##### Experimental Model and Subject Details

All strains were grown with aeration in Luria-Bertani (LB-Miller, BD-Difco) broth at 37°C except for the *E. coli* lambda cI857 lysogen which was grown at 30°C. Strains used in this study are listed in Table S1. Unless otherwise noted, antibiotics and inducers were used at: 100 µg $mL^{-1}$ ampicillin (Amp, Sigma), 100 µg $mL^{-1}$ kanamycin (Kan, GoldBio), 5 µg $mL^{-1}$ chloramphenicol (Cm, Sigma), 100 ng $mL^{-1}$ anhydrotetracycline (aTc, Clontech), and 0.4 mM Isopropyl β-D-1-thiogalactopyranoside (IPTG, GoldBio). For microscopy, HALO-TMR (Promega) and SNAP-$JF_{503}$ (Lavis Lab) were used at concentrations of 1 and 2 µM, respectively. AB solid agar consisted of 1.5% agar, 0.3 M NaCl, 50 mM $MgSO_4$, 0.2% casamino acids, 1 mM arginine, 1% glycerol, and 10 mM potassium phosphate at pH 7.5.

##### Cloning Techniques

Primers and dsDNA (gene blocks) used for plasmid construction are listed in Table S2, and all were obtained from Integrated DNA Technologies. Plasmids are listed in Table S3. Plasmid combinations are organized according to the figures and strains in which they appear in Table S4. Gibson assembly, intramolecular reclosure, and traditional cloning methods were employed for all cloning, as indicated in Table S2. PCR with Q5 High Fidelity Polymerase (NEB) was used to generate insert and backbone DNA. Gibson assembly relied on the HiFi DNA assembly mix (NEB). All enzymes used in cloning were obtained from NEB. Construction of the random $cI_{VP882}$-HALO variants was carried out by GeneMorph II EZClone (Agilent) using 250 ng of template DNA (pJES-183), primers JSO-1649/1650, and 32 cycles. Transfer of plasmids into the *V. parahaemolyticus* VP882 lysogen was carried out by conjugation followed by selective plating on 50 U $mL^{-1}$ polymyxin B (Sigma) and Kan. In *V. parahaemolyticus*, 10 ng $mL^{-1}$ aTc was used for induction rather than the 100 ng $mL^{-1}$ inducer concentration used in *E. coli*.

##### Growth, Lysis, and Reporter Assays

Unless otherwise noted, overnight cultures were back diluted 1:200 into fresh medium with appropriate antibiotics prior to being dispensed (200 µL) into 96 well plates (Corning Costar 3904). aTc was added as specified. Plates were shaken at 37°C or 30°C and a BioTek Synergy Neo2 Multi-Mode reader was used to measure $OD_{600}$ and bioluminescence. Relative light units (RLU) were calculated by dividing the bioluminescence by the $OD_{600}$ at that time. Testing the panel of SNAP-Qtip mutants (Figure 4B) made additional use of a BioSpa 8 automated incubator to read multiple plates at regular intervals. Unless otherwise noted, fixed time-point reporter assays were measured ~8 h after inoculation of the cells into the wells.

**Protein Expression and Purification**

For production and purification of $cI_{Lambda}$-HALO-HIS, $cI_{VP882}$-HALO-HIS, the chimeras, and $cI_{VP882}$-HALO-HIS variants used for microscopy, cleavage assays, and DNA-binding (P*q-lux*) assays, plasmids harboring the indicated constructs were expressed in *E. coli* T7Express lysY/$I^q$ using 0.4 mM IPTG induction and 18°C incubation overnight. Cells were pelleted at 16,100 x g for 10 min and resuspended in lysis buffer (25 mM Tris-HCl pH 8, 150 mM NaCl, 1 mM DTT). Cells were lysed using sonication and subjected to centrifugation at 32,000 x g for 40 min. The supernatants were applied to Ni-NTA Superflow resin (Qiagen). Resin was washed with 5 column volumes (CVs) of 25 mM Tris-HCl pH 8, 150 mM NaCl, 20 mM imidazole, and 1 mM DTT, and eluted in 2 CVs of 25 mM Tris-HCl pH 8, 150 mM NaCl, 300 mM imidazole, and 1 mM DTT. Eluates were concentrated, incubated with 7 µM (final conc.) HALO-$Alexa_{660}$ ligand on ice for 30 min before being loaded onto a Superdex-200 size exclusion column (GE Healthcare) in gel filtration buffer (25 mM Tris-HCl pH 8, 150 mM NaCl, and 1 mM DTT). For differential labeling experiments requiring the use of both HALO-$Alexa_{660}$ and HALO-TMR labeled WT $cI_{VP882}$-HALO, $cI_{VP882}^{S130A}$-HALO, and $cI_{VP882}^{K172A}$-HALO proteins, a second batch of lysates prepared from the relevant strains was purified using the identical procedure as for the HALO-$Alexa_{660}$ labeled proteins except that the label used was HALO-TMR (5 µM final conc.). All proteins were concentrated, flash frozen, and stored at -80°C prior to use. Differentially labeled proteins were stored separately and mixed immediately before use.

For production and purification of $cI_{VP882}$-HALO (no HIS) used for comparative analysis to $cI_{VP882}$-HALO in complex with HIS-Qtip, $cI_{VP882}$-HALO (no HIS) (pJES-190) was overexpressed in *E. coli* T7Express lysY/$I^q$ as above. The clarified lysate was loaded onto a heparin column (GE Healthcare) and eluted by a linear gradient from buffer A (25 mM Tris-HCl pH 8, 1 mM DTT) to buffer B (25 mM Tris-HCl pH 8, 1 M NaCl, 1 mM DTT). Peak fractions were pooled, concentrated, treated with 7 µM HALO-$Alexa_{660}$ ligand (final conc.), subjected to Superdex-200 size exclusion chromatography in gel filtration buffer, and stored at -80°C.

Production and purification of HIS-Qtip in complex with $cI_{VP882}$-HALO was carried out as above with the HIS-tagged proteins, except that the *E. coli* T7Express lysY/$I^q$ co-expressed HIS-Qtip (pJES-147) and $cI_{VP882}$-HALO (pJES-190). The eluted complex was concentrated, treated with 7 µM HALO-$Alexa_{660}$ ligand (final conc.), subjected to Superdex-200 size exclusion chromatography in gel filtration buffer, and stored at -80°C.

**Electrophoretic Mobility Shift Assay (EMSA)**

Primers JSO-956/957 (Table S2) and purified VP882 DNA were used to make the probe (257 bp). 20 ng of probe was used in each EMSA reaction. The highest concentration of $cI_{VP882}$-HALO, designated 25x (Figure 1B), was ~300 nM. The concentration of the HIS-Qtip-$cI_{VP882}$-HALO complex was matched to that of free $cI_{VP882}$-HALO (no HIS-Qtip) based on the fluorescence from the $cI_{VP882}$-HALO-conjuaged $Alexa_{660}$ dye present in both samples. A 5-fold serial dilution of protein was applied to indicated lanes. The protein and probe were combined in binding buffer (25 mM Tris-HCl pH 8, 50 mM NaCl, 1 mM DTT), and incubated at RT for 15 min. The samples were subsequently loaded onto a Novex 6% DNA Retardation Gel (Thermo) at 4°C and electrophoresed in 1x TBE at 130 V for 2 h. The gel was stained for DNA by SYBR Green I Nucleic Acid Gel Stain (Thermo) and washed in 1x TBE for 20 min. The gel was subsequently imaged using an ImageQuant LAS 4000 imager under the SYBR Green setting. A duplicate batch of the samples was also subjected to electrophoresis in a 4-20% SDS-PAGE gel to verify the correct loading of the HALO-$Alexa_{660}$-labeled $cI_{VP882}$-HALO in the EMSA. The gel was imaged using an ImageQuant LAS 4000 and detection of HALO and SNAP as outlined below.

***in vitro* Cleavage of Phage Repressor Proteins**

10 μL of the indicated HALO-tagged repressor protein (0.5 mg $mL^{-1}$) was mixed with 0.5 μL JSO-1538 ssDNA (1 mM, Table S2), 1 μL ATP-γ-S (50 mM, Sigma), and 10 μL 10x RecA buffer (NEB) in a 100 μL preparation. 2.5 μL RecA (2 mg $mL^{-1}$, NEB) was added to initiate the reaction. All cleavage reactions were carried out at 37°C. At each time point, 10 μL aliquots were taken from the reactions and mixed with 5 μL 4x Laemmli Sample Buffer (BioRad). The samples were loaded onto a 4-20% SDS-PAGE gel for electrophoresis. The gel was imaged using an ImageQuant LAS 4000 imager under the Cy5 or the Cy3 and Cy5 settings before being stained with Coomassie Brilliant Blue. The RecA band revealed by Coomassie was used as the loading control (see below for additional details on in-gel imaging). In Figure 2A, RecA and ATP-γ-S were individually withheld from the reactions, as indicated in the figure. For the mixed reactions (Figure 3A), the total amount of $cI_{VP882}$-HALO protein was doubled (20 μL per 100 μL reaction). Due to the extremely low yield of soluble HIS-Qtip-$cI_{VP882}$-HALO complex that could be obtained using our approach, in Figure 2B, approximately 20-fold less $cI_{VP882}$-HALO protein was used in the reactions relative to those in the other cleavage assays reported here. To ensure that equal amounts of free $cI_{VP882}$-HALO and $cI_{VP882}$-HALO in complex with HIS-Qtip were loaded onto gels,

the amount of free $cI_{VP882}$-HALO, as judged by HALO-$Alexa_{660}$ fluorescence, was adjusted to match that of the labeled $cI_{VP882}$-HALO in complex with HIS-Qtip.

**In-Gel (Western) Detection of HALO and SNAP Constructs**

SDS-PAGE gels were imaged for labeled HALO. HALO-TMR (excitation/emission: 555/585 nm) and HALO-$Alexa_{660}$ (excitation/emission: 663/690 nm) were detected in a LAS 4000 imager under the Cy3 and Cy5 settings, respectively. Gels were subsequently stained for total protein using Coomassie Brilliant Blue. For assaying SNAP-Qtip production levels (Figure S4A), overnight cultures of the indicated strains were back diluted 1:500 in fresh medium containing 100 ng $mL^{-1}$ aTc, 1 µM HALO-TMR, and 2 µM SNAP-Cell 647-SiR (NEB). Cultures were grown for 8 h at 37°C, prior to cell harvest, lysis (BugBuster, Millipore), and electrophoresis by SDS-PAGE. SNAP-Cell 647-SiR (excitation/emission: 647/661 nm) was detected by the LAS 4000 imager under the Cy5 setting making it compatible to use with HALO-TMR. Exposure times for all fluorophores never exceeded 30 sec.

**Confocal Microscopy**

Overnight cultures of *E. coli* T7Express lysY/$I^q$ carrying the indicated HALO and/or SNAP fusions were diluted 1:500 in medium containing 100 ng $mL^{-1}$ aTc, 1 µM HALO-TMR, and/or 2 µM SNAP-$JF_{503}$. Cultures were returned to growth for 4-6 h, subjected to centrifugation, washed with PBS, and resuspended in fresh PBS to a final $OD_{600}$ = 0.1-0.3. 8 µL of each sample was spotted onto a glass coverslip and overlaid with an AB agar pad. Samples were imaged approximately 40 min after preparation on a Leica SP8 Confocal microscope under photon counting mode. HALO-TMR was excited with 561 nm light and detected within the range 569-625 nm. SNAP was excited at 499 nm and detected between 513-542 nm. When imaged together, the HALO and SNAP channels were collected in sequential mode. Images for each sample were acquired as a tile-scan (3x3 or 4x4) and stitched in LASX (Leica Microsystems).

**Image Analysis**

Image analyses were performed in FIJI software (Version 1.52p). To generate intensity line traces and composite images of fluorescent SNAP and HALO fusions, individual cells were first segmented in the brightfield channel via an intensity-based thresholding procedure after image smoothing. Only cells of a defined size range were retained in further analysis for consistency. Individual segmented cells were manually rotated using the rotated rectangle function such that the long axis of the cell was oriented in the x-dimension. In cases where one cell poll contained a punctate signal, this pole was aligned to the left side of the image. For each x-aligned cell, the intensity profile was measured through a 0.5 µm thick (y-dimension) line traced through the long axis of the cell. Data were exported for quantitation and graphing in R software using gglplot2 (https://ggplot2.tidyverse.org). To account for cell-to-cell intensity differences, each line trace was normalized to the peak pixel intensity for that cell. For each condition (strain, fluorophore), the intensity traces of the individual cells were averaged, and the first standard deviation calculated to generate plots. To produce a composite image for each condition, the aligned individual cell images were concatenated and subsequently averaged. Resulting composite images were normalized to the highest pixel value within that image. For display, enhanced lookup tables were used (Red Hot for HALO-TMR fusions and Cyan Hot for SNAP-$JF_{503}$ fusions). In all cases, n > 20 cells were analyzed.

##### Quantification and Statistical Analysis

Data are presented as the mean ± std unless otherwise indicated in the figure legends. The number of independent biological replicates for each experiment is indicated in the figure legends or methods. For imaging analysis, 20-25 single cells were used in compiling each composite. Cells were excluded from analysis if they were abnormal in size, contacting another cell, and/or exhibiting poor fluorescence from the SNAP- or HALO- channel.

##### Data and Software Availability

Unprocessed gels and micrographs that were used to in this study are deposited on Zenodo (10.5281/zenodo.3576390). Other experimental data that support the findings of this study are available by reasonable request from the corresponding author.

### Supplemental Tables

**Table S1, related to Methods. Bacterial strains used in this study.**

(Available as a separate Excel file)

**Table S2, related to Methods. Oligonucleotides and gene blocks used in this study.**

(Available as a separate Excel file)

**Table S3, related to Methods. Plasmids used in this study.**

(Available as a separate Excel file)

**Table S4, related to Methods. Plasmids used in each figure in this study.**

(Available as a separate Excel file)
